# Near Infrared-II Fluorescent protein for In-vivo Imaging

**DOI:** 10.1101/2022.03.04.482971

**Authors:** Zong Chang, ChenChen Liu, Shubi Zhao, Jiaqi Chen, Xiaoping Zhang, Hanyu Tian, Qinchao Sun

## Abstract

In vivo fluorescent imaging in the second near-infrared window (NIR-II) provides an excellent approach for understanding the biological processes in substantially scattered tissue environments with reasonable temporal-spatial resolution. In spite of an enormous amount of organic and inorganic NIR-II fluorophores developed, there is no NIR-II fluorescent protein reported. Here, we present the first NIR-II fluorescent protein, IRFP1032 which exhibits strong exciton absorption and emission in the NIR-II region, with exciton extinction coefficient about 4.1 ×10^6^ M^-1^cm^-1^ at the excitation maximum 1008 nm, emission maximum of 1032 nm, and emission quantum yield about 0.84%. The IRFP1032 is found to be the brightest NIR-II fluorophore ever reported (brightness of 3.4 × 10^4^ M^-1^cm^-1^ in PBS) which is thousands-fold brighter than IR26 in DCM. Taking the advantage of the excellent photo-properties of the NIR-II fluorescent proteins, a collection of high-quality in vivo imaging research was realized, for instance, real time observation of blood flow dynamics, dual-channel imaging of the lymphatic/blood vessel network and the trajectories of single bacterial cell travelling in blood vessels. Moreover, a mammalian expression vector was constructed for the IRFP1032, and the corresponding NIR-II fluorescence was able to be recorded unambiguously. The promising NIR-II in vivo imaging properties of IRPF1032 demonstrated here would open a new scene in fluorescent protein-based imaging.

## Introduction

In vivo fluorescence imaging has been significantly advancing the understanding of fundamental biology and clinical medicine as the capability of achieving high resolution images in space and time.^1–3^ The optimal photon energy for in vivo imaging is considered to be located in the near infrared window (NIR, 700 nm – 1700 nm), especially in the second near infrared window (NIR-II, 1000 nm – 1700 nm), since in which region the photon scattering, photon absorption as well as autofluorescence of tissue are reasonably compromised.^4–8^ The NIR fluorophores as the contrast probe are the critical to the performance of in vivo fluorescence imaging.^9,10^ The higher absorption and emission efficiency (brightness), along with lower emission energy of fluorophores (NIR-II region), the more enhanced in vivo image quality could be realized.^10^ Thanks to the development of material science and biological technology, an enormous variety of near infrared fluorophores have been designed for in vivo imaging over the last decades.^11,12^

The first high contrast in vivo intact blood vessel fluorescence imaging was reported by Dai et al., via intravenous injection of the surface modified NIR-II emitting single wall carbon nanotubes (utilizing the emission peaks above 1000 nm).^13,14^ Following their pioneer work, motivated by the great potential of NIR-II imaging in the clinical application, a tremendous amount of fluorophores have been developed with emission in the NIR-II region, such as the semiconductor quantum dots with the right bandgap in the NIR-II region (characterized by the high NIR-II emission quantum yield),^15,16^ the lanthanide doped down conversion nanoparticles (sharp NIR-II emission bands and tunable emission lifetime),^17,18^ the organic molecules with large π-conjugation system (relative lower cytotoxicity and higher brightness),^19–21^ etc.^22–24^

Most of the reported NIR-II fluorophores were able to demonstrate high contrast in vivo fluorescence imaging for blood vessels and tumor tissue.^16,25^ Nevertheless, the poor cell and tissue specific targeting of current NIR-II fluorophores greatly limits the NIR-II imaging as the powerful platform for in vivo biological study, for instance immune cell tracking, stem cell distribution in vivo observation and neuron imaging. In contrast, the genetically encoded fluorescent protein could be precisely expressed in the targeted cells or even in specific organelles such as cell membrane, mitochondria, etc.^26–28^ However, unlike the organic and inorganic fluorophores for which the emitted photon energy could be flexibly adjusted via varying the functional groups, size or element composition, etc.,^15,17,22^ the measures to extend the emission spectrum of genetically encoded fluorescent protein are rather constrained by the protein structure.^29,30^ For example, in order to extend the emission wavelength of the most famous beta-barrel shaped fluorescent protein (e.g. Green Fluorescence Protein), great efforts have been made to modify the protein geometry via gene mutation, yet, as far as we know, there is no NIR fluorescent protein successfully developed based on such geometry.^30^ Besides the beta-barrel shaped fluorescence protein, another kind of fluorescence protein, which is constituted by the polypeptide units and the fluorophore ligand, exhibits strong fluorescence emission via the unique interaction between the two components (the emission of individual ligand not observable or well shifted from the protein complex emission).^31–36^ In the meanwhile, such kind of fluorescent protein provides a more diverse way for extending the emission spectrum. Tsien et al. have reported a novel fluorescent protein (IFP1.4) which was found from bacterial phytochrome and exhibits strong NIR emission with maximum around 708 nm via binding with biliverdin.^35^ Intensive studies have been conducted on such fluorescent protein system to improve the photo-properties and to functionalize as a calcium indicator for neuron communication.^32–34^

Actually, there is a huge family of protein complex with excellent photo-properties, the so-called light harvesting complex, for example, the B-Phycoerythrin from the light harvesting complex of certain red algae is considered to be the brightest fluorophore with quantum yield about 0.98, absorption around 566 nm (similar to the individual ligand) and emission around 576 nm,^31^ which has been widely used (commercialized by Thermo Fisher) in immunofluorescent staining, flow cytometry, etc. Unfortunately, the emission wavelength of the reported fluorescence proteins is still located in the relatively high scattering and autofluorescence region for in vivo imaging. To discover novel fluorescence proteins with emission wavelength in the NIR-II region would substantially promote the fundamental research on biology via in vivo fluorescence imaging.

As is well-known, the purple photosynthetic bacteria like other photosynthetic species are able to utilize photon energy for biosynthesis, while the photon energy collected by purple photosynthetic bacteria is much lower than that by the red algae and green plants.^37,38^ Various kinds of light harvesting protein of purple photosynthetic bacteria have been found with the capability of absorbing the solar energy in NIR window.^37,38^ In general, the light harvesting protein acts as antenna to receive the photon energy and resonantly transfer the energy to the reaction center for biosynthesis.^38^ Besides the efficient energy transfer from the excited state of light harvesting protein, the thermal quenching process of the excited state of light harvesting protein might be extremely suppressed.^38^ Therefore, we might expect a reasonable amount of the radiative transition from the excited state of the light harvesting protein, even while the reaction center is involved. Herein, we carefully studied two kinds of light harvesting complexes and found the first NIR-II fluorescence proteins IRFP1032 which exhibits strong exciton absorption and emission in NIR-II region and is found to be the brightest NIR-II fluorophore ever reported. The IRFP898 as another NIR fluorescent protein with well separated emission bands was also reported. A collection of fluorescence imaging experiments was performed to demonstrate the brilliant capability of the IRFP1032 and IRFP898 for in vivo biological research, for instance blood vessels imaging, blood flow dynamics, dual color imaging, and single bacterial cell tracking. Furthermore, a mammalian expression vector was constructed for the IRFP1032, and the corresponding NIR-II fluorescence was able to be observed unambiguously.

## Results and Discussion

### Fluorescent protein preparation

The light harvesting protein as a kind of membrane protein was isolated according to the standard procedure with a few modifications (Figure S1a).^39^ Briefly, the harvested bacteria were crushed and the fluorescence protein membrane was collected via ultra-centrifugation. 3% (w/w) *n*-dodecyl β-d-maltoside (DDM) was used to dissolve the fluorescence protein from membrane and a sucrose step gradient ultra-centrifugation was followed. The resulting fluorescence protein was isolated from the sucrose gradient according to the exciton absorption band and then washed through a 100KD cut off centrifuge filter to remove the excess DDM. Furthermore, the resulting fluorescence protein was able to pass through a 300KD cut off centrifuge filter also indicating that the fluorescence protein was well dissolved from the membrane. During the purification process, the reaction centre protein as enveloped in the light harvesting protein ring was isolated along with the fluorescence protein. The purity of the isolated fluorescence protein was characterized by the ratio of the absorbance at exciton absorption maximum to that at 280 nm, as reported by Neil Hunter et al. for Cryo-EM experiment.^39^ For IRFP1032, the ratio of absorbance at 1008 nm to that at 280 nm was around 1.15, and for IRFP898, the ratio of absorbance at 878 nm to that at 280 nm was around 1.2, as shown in Figure S1b, S1c, which are quite close to the reported purity for Cryo-EM.^39^ HPLC (on AKTA) was applied to further characterize the purity of the isolated proteins by elution of 5x PBS solution 0.5 mL/min as shown in Figure 1Sa. The retention time was found with about 15 to 20 mins both IRFP1032 and IRFP898, with no other impurities observed. The exciton absorption spectrum of the isolated fluorescence protein (IRFP1032, IRFP898) was in reasonable agreement with the reported absorption of light harvesting protein from the same purple photosynthetic bacteria species,^38,39^ detailed discussion in the following section.

**Figure 1.**
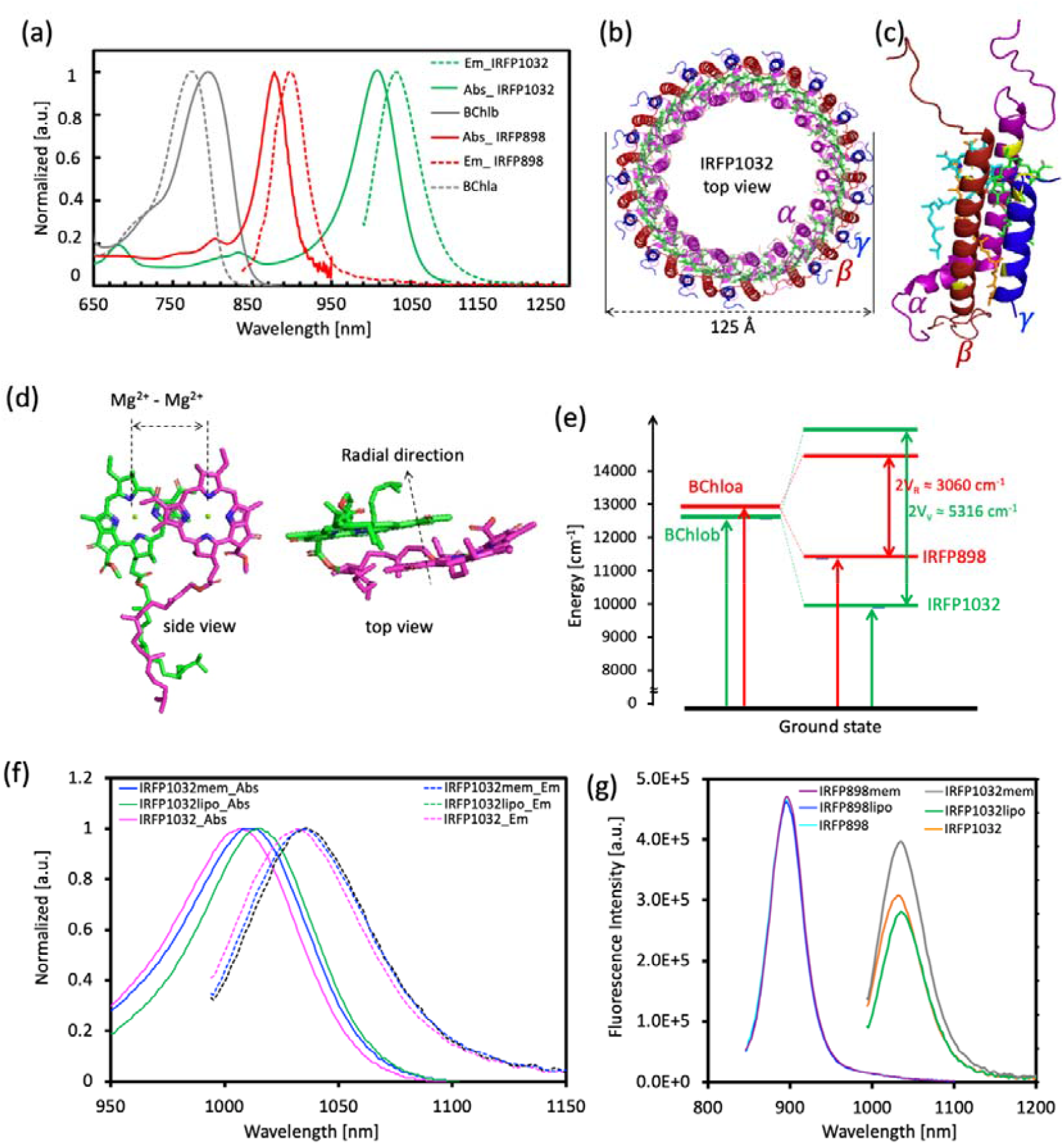
The photophysical properties and protein structure of IRFP1032 andIRFP898. (a) The absorption spectrum of the bacterial chlorophyll a (BChla) in methanol (grey dash line), the absorption spectrum of bacterial chlorophyll b (BChlb) in methanol (grey solid line), the absorption and emission spectra of IRFP1032 in PBS (green lines) and the absorption and emission spectra of IRFP898 in PBS (red lines). (b) The top view of the 3D structure of IRFP1032, the magenta helix (*α* subunit), the red helix (*β* subunit), the blue helix (*γ* subunit) and the complex ligands (green), PDB:6ET5. (c) The interaction between one of the BChlb dimers and *α, β, γ* subunits, the yellow sites represent the histidine for *α, β* subunit and the tryptophan in *γ* subunit (the corresponding amino acid sequence highlighted in red in Figure S1f). (d) The top and side view of the BChlb dimer in the IRFP1032. (e) The energy levels of the BChla, BChlb, the IRFP1032 and IRFP898. The V_R_ represents the absorption energy difference between BChla and IRFP898 (coupling energy), Vv represents the absorption energy difference between BChlb and IRFP1032 (coupling energy). (f) The normalized absorption and emission spectra of IRFP1032 in different environments, the IRFP1032mem as in cell membrane (before isolation), IRFP1032 as isolated from membrane by DDM, the IRFP1032lipo as in liposome. (g) The relative emission intensity of IRFP1032 and IRFP898 in different environments (in cell membrane, liposome, and isolated with DDM) with the same absorbance at the excitation wavelength, for IRFP1032 at 980 nm and for IRFP898 at 825 nm.

### Electronic and structural properties of IRFP1032 and IRFP898

The photo-properties of the IRFP1032 and IRFP898 are depicted in Figure 1a. The excitation energy of the IRFP1032 is located in the NIR-II region with absorption maximum about 1008 nm which is the lowest electronic excitation energy for protein complexes have ever been reported.^8^ The absorption band of IRFP1032 in the NIR-II region is featured as narrow peak (full width half maximum (FWHM) about 60 nm) and without fine structure. In comparison to IRFP1032, the absorption band of IRFP898 is significantly blue shifted by 130 nm, with maximum about 878 nm in the NIR-I region and FWHM around 50 nm. A relatively strong NIR-II emission was observed for IRFP1032 with a maximum around 1032 nm. The small Stokes shift (24 nm) between the emission and absorption band of IRFP1032 indicates the quite “rigid” geometry of the ground and excited state. A similar Stokes shift (20 nm) was found for IRFP898 with emission maximum about 898 nm. The absorption band of the bacteriochlorophyll b (BChlb, ligand of IRFP1032) is around 796 nm in methanol, which is about 212 nm blue shifted to the absorption of IRFP1032. The absorption band of bacteriochlorophyll a (BChla, ligand of IRFP898) is about 771 nm and 107 nm blue-shifted to that of IRFP898. The lower electronic absorption energy observed for BChlb might be caused by the expanded π-conjugation chain, for which an additional C=C double bond could be found, instead of the saturated bond in BChla, as indicated by the red circle in Figure S1d, S1e. The large red shifted excitation energy for the protein complex (IRFP1032, IRFP898) to the individual chlorophyll (BChla, BChlb) would be crucial in the case of imaging via in vivo expression of such fluorescent protein complex as which extremely suppresses the interference from the fluorescence of the unbinding ligands during excitation.

The dramatic red-shifted excitation energy of the fluorescent protein to that of individual bacterial chlorophyll might arise from the unique protein complex architecture, which render a strong inter-chlorophyll electronic coupling and significant protein-chlorophyll interaction.^38^ The 3D geometry of the IRFP1032 could be mapped by the cryo-EM, as shown in Figure 1b with 2.9 Å resolution.^39^ There are 17 *α* and *β* helix subunits, 16 *γ* helix subunits and 34 bacteriochlorophyll b to construct the well-known ring shape of the IRFP1032. In spite of the 3D geometry of the IRFP898 has not been reported, a ring shape of IRFP898 with the 16 *α*, 16 *β* helix subunits and 32 bacteriochlorophyll a could still be demonstrated by cryo-EM with 8.5 Å resolution.^40^ As revealed by the cryo-EM, the bacteriochlorophylls of IRFP1032 are arranged in the head to tail structure to form the so-called J aggregation,^38,41–43^ Figure 1b, 1d. The strong electronic coupling between the J aggregated chlorophylls renders the individual chlorophylls to act coherently as a single molecule. Therefore, while interacting with photons, the excited states of the IRFP1032 are not localized on certain individual chlorophyll anymore, instead, all the chlorophylls in the protein complex coherently share the same excited electron, which also known as exciton. The electronic coupling effect splits the electronic states of the protein complex (IRFP1032, IRFP898) into a higher energy state and a lower energy state, as shown in Figure 1e. The energy difference between the higher state and the lower state is known as the splitting energy which reveals the strength of the electronic coupling effect. The splitting energy is calculated by twice over the absorption peak energy difference between the protein complex and the corresponding individual bacteriochlorophyll which is about 3060 cm^-1^ and 5316 cm^-1^ for IRFP898 and IRFP1032, respectively. The much higher splitting energy of IRFP1032, which is about 1.7 times larger than that of IRFP898 indicates a much stronger electronic coupling between the bacteriochlorophylls. One of the main factors that affects the electronic coupling is considered to be the distance between each chlorophyll (Mg^2+^ to Mg^2+^ distance, Figure 1d). However, by investigating the external diameter of the ring of IRFP1032 (about 125 Å^39^) and IRFP898 (about 115 Å^40^), we found that the Mg^2+^ to Mg^2+^ distance of both protein complex is almost the same about 11.5 Å (assuming the bacterial chlorophylls equally spaced on the ring, 32 chlorophylls for IRFP898 and 34 chlorophylls for IRFP1032). Therefore, the strong coupling of the IRFP1032 might be mainly caused by the extra *γ* helix, for which the rich of phenyl group and hydrogen bond site might push the two-chlorophyll plane much closer along the radial direction than that of IRFP898, Figure 1d.

The extra *γ* helix of IRFP1032 would render a much sensitive response to the surrounding environment on the absorption and emission properties, as shown in Figure 1f, 1g, S1g. For IRFP898 with absence of *γ* helix, there is no significant changes observed in the exciton absorption, emission energy and intensity under different environments, Figure S1g. While for IRFP1032, a significant emission enhancement was observed in the membrane state (IRFP1032mem) than that in the liposome (IRFP1032lipo) and isolated with DDM (IRFP1032), Figure 1g. The exciton absorption and emission maximum were found to be slightly red shifted for the IRFP1032 in the membrane and liposome state to DDM isolation state.

### Exciton brightness of IRFP1032 and IRFP898

The brightness (EC*QY) is one of the most important parameters of fluorophores for the success of in vivo imaging.^8,10^ However, as the emission wavelength getting longer (from NIR-I to NIR-II), the nonradiative decay becomes more and more critical in the excited state relaxation path, especially for organic fluorophores.^22^ In order to synthesize organic dyes with NIR-II emission, an extended π-conjugation chain is the most commonly applied molecular design strategy.^11,12^ The large π-conjugation system, on one hand, significantly reduces the energy gap between the excited state and ground state resulting a long wavelength emission (NIR-II); on the other hand, the degrees of freedom of such system is dramatically increased, as a consequence the photoinduced isomerization, bond rotation and charge transfer would severely quench the excited state. As far as we known, most of the reported NIR-II organic dyes exhibits very low quantum yields.^11,12,22^ Even worse, the fluorescence quantum yield (QY) and extinction coefficient (EC) of most NIR-II organic dyes is further significantly reduced in aqueous solution.^12,19^ The extinction coefficient, emission spectra and the emission quantum yield of IRFP1032 and IRFP898 are shown in Figure 2 and Table 1. A strong NIR-II emission could be observed from the IRFP1032 in comparison to the commercial NIR-II dye IR26 with QY about 0.84%, which is about 28-fold higher than that of IR26 in DCM, in Figure 2a (the blue and orange solid curve). The emission quantum yield of IRFP898 was found to be as high as 1.9% as reference to the IR820 in EtOH. Besides the high emission quantum yields, the extinction coefficient of the exciton absorption of IRFP1032 and IRFP898 was also found to be much higher than that of most organic dyes due to the significant electronic coupling effect, 4.1 × 10^6^ M^-1^cm^-1^ and 3.4 × 10^6^ M^-1^cm^-1^ at the exciton absorption maximum for IRFP1032 and IRFP898, respectively (similar order of magnitude of B-Phycoerythrin^47^). A few recently reported organic dyes (absorption and emission maximum in or very close to NIR-II region) with different molecular design strategies are summarized in Table 1 and Figure S2. Thanks to the strong exciton extinction coefficient and the high NIR-II emission quantum yield, to the best of our knowledge, the IRFP1032 is found to be the brightest NIR-II fluorophore that ever been reported (brightness about 3.4 × 10^4^ M^-1^cm^-1^), about 3-orders of magnitude higher than IR-26 in DCM.

**Table 1.** The photophysical properties of several NIR-II fluorophores

| Fluorophore | Abs<sub>max</sub> (nm) | Em<sub>max</sub> (nm) | EC<sub>max</sub> (M<sup>-1</sup>cm<sup>-1</sup>) | QY (%) | Brightness<sub>max</sub> (M<sup>-1</sup>cm<sup>-1</sup>) | Solvent |
| --- | --- | --- | --- | --- | --- | --- |
| IRFP1032<sup>a</sup> | 1008 | 1032 | 4,100,000 | 0.84 | 34,000 | PBS |
| IRFP898<sup>a</sup> | 878 | 898 | 3,400,000 | 1.9 | 65,000 | PBS |
| LZ1105<sup>b</sup> | 1041 | 1105 | 100,000 | 0.03 | 30 | PBS |
| LZ1105<sup>c</sup> | 1041 | 1105 | 150,000 | 3.9 | 5,800 | DMSO |
| Flav7(3)<sup>d</sup> | 1026 | 1045 | 240,000 | 0.3 | 720 | DCM |
| FM1210<sup>e</sup> | 980 | 1210 | 27,000 | 0.03 | 8.1 | DCM |
| IR26<sup>f</sup> | 1073 | 1138 | 110,000 | 0.03 | 33 | DCM |
| IR820<sup>g</sup> | 820 | 850 | 200,000 | 4.4 | 8,800 | EtOH |

<sup>a</sup>QY of IRFP1032 about 0.84 ± 0.15% to IR26 in DCM, QY of IRFP898 about 1.9 ± 0.07% to IR820 in EtOH, <sup>b</sup>reported as QY=0.03% to IR26 in DCE (0.05%), <sup>19</sup><sup>c</sup>reported as QY=3.9% to IR26 in DCE (0.05%), <sup>19</sup><sup>d</sup>reported as QY=0.5% to IR26 in DCM (0.05%), <sup>21</sup><sup>e</sup>reported as QY=0.03% to IR26 in DCE (0.05%), <sup>24</sup><sup>f</sup>IR26 in DCM with QY=0.03%, <sup>44,45</sup><sup>g</sup>IR820. <sup>46</sup>

**Figure 2.**
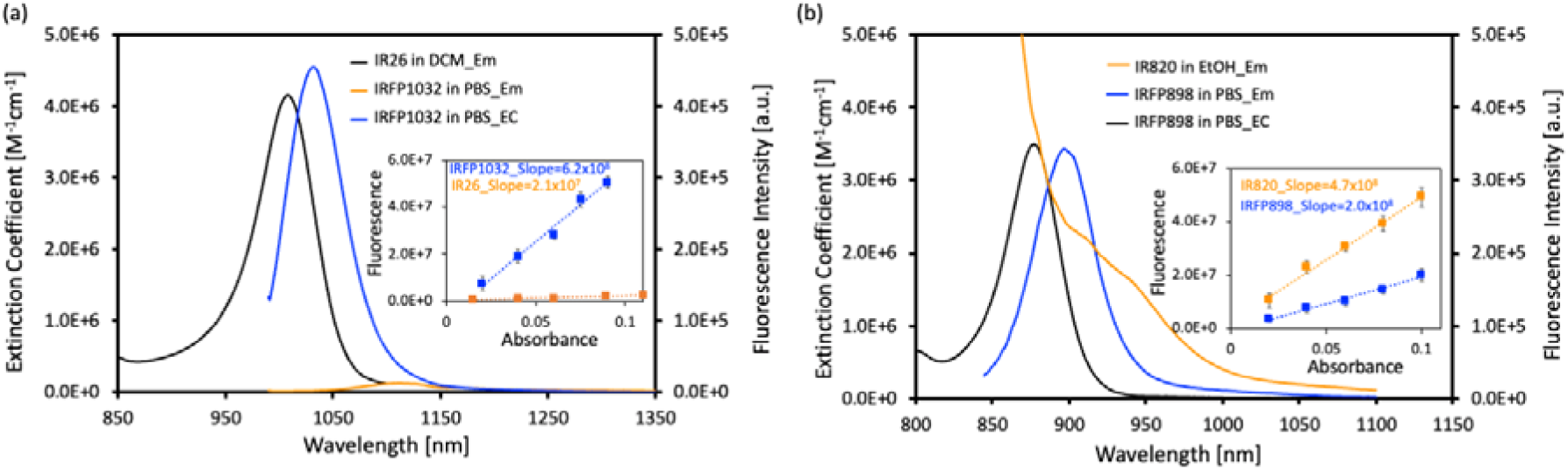
The exciton brightness characterization of IRFP1032 and IRFP898 in PBS. (a) The extinction coefficient (EC) of IRFP1032 (black); the emission spectrum of IRFP1032 in PBS (blue) and IR26 in DCM (orange) with the same absorbance at 980 nm around 0.1 (optical path 1 cm). Inset: the integrated emission intensity from 1000 nm to 1500 nm as a function of the absorbance at 980 nm, the blue solid square for the IRFP1032 in PBS and orange solid square for IR26 in DCM, the dash line is the linear fitting of the corresponding data. (b) The extinction coefficient (EC) of IRFP898 (black); the emission spectrum of IRFP898 in PBS (blue) and IR820 in EtOH (orange) with the same absorbance at 825 nm around 0.1 (optical path 1 cm). Inset: the integrated emission intensity from 850 nm to 1200 nm as a function of the absorbance at 825 nm, the blue solid square for the IRFP898 in PBS and orange solid square for IR820 in EtOH, the dash line is the linear fitting of the corresponding data.

### In vivo imaging with IRPF1032 and IRFP898

The visible and NIR photons would suffer from severe scattering while travelling through the tissue environment, however as the photon energy getting lower (in NIR-II), the corresponding tissue scattering effect could be dramatically reduced (the wavelength dependence of the scattering is the combination of Rayleigh scattering and Mie scattering, also tissue dependent).^6–8^ In spite of the scattering effect dramatically reduced in the NIR-II region, the scattering background still acts as a dominant factor on the contrast of in vivo imaging (Figure S3, Table S1).^45^ As shown in Figure 3a, the in vivo fluorescence imaging of mouse hindlimb blood vessels was recorded with long pass filters from 1200 nm to 1400 nm. The IRFP1032 was injected in the form of liposome embedded to prolong the blood circulation time (Figure S15). As illustrated in Figure 3b, S5b, the ratio of the hindlimb vessel fluorescence intensity to the adjacent tissue background (SBR^a^) was slightly increased by switching the long pass filter from 1200 nm LP (SBR^a^ 1.1) to 1300 nm LP (SBR^a^ 1.2). However, significant enhancement on the image contrast could be observed for over 1400 nm imaging with SBR^a^ about 1.9 (Figure 3b, S5b). Regarding to the imaging contrast, the well-known ballistic photon (non-scattered photon) is considered to be the only effective photon for sharp imaging, while the scattered photon makes image blurry considered to be scattering background.^8^ Besides the reduced scattering effect, the remarkably reduced background for over 1400 nm in vivo imaging might be also attributed to the strong absorption of scattered photons by water, since the water overtone absorbance of light around 1400 nm is about 10 times higher than that of 1200 nm and 1300 nm (Figure S7b).^48^ A similar enhancement on the image quality could also be observed for through intact scalp and skull imaging as shown in Figure 3c, S5c, as well as the scalp removed brain vessels imaging in Figure S4, S5d. A brain vessel with FWHM about 73 μm could be clearly resolved through intact scalp and skull for over 1400 nm imaging, yet no clear vessel structure could be observed at the same location in the case of over 1300 nm imaging (Figure 3c, S6c). Furthermore, brain capillaries were clearly imaged as shown in Figure S14. As demonstrated, the fluorescent protein of IRFP1032 exhibits outstanding capability of NIR-II in vivo imaging from 1200 nm to 1400 nm, especially in the high contrast imaging region of over 1400 nm.

**Figure 3.**
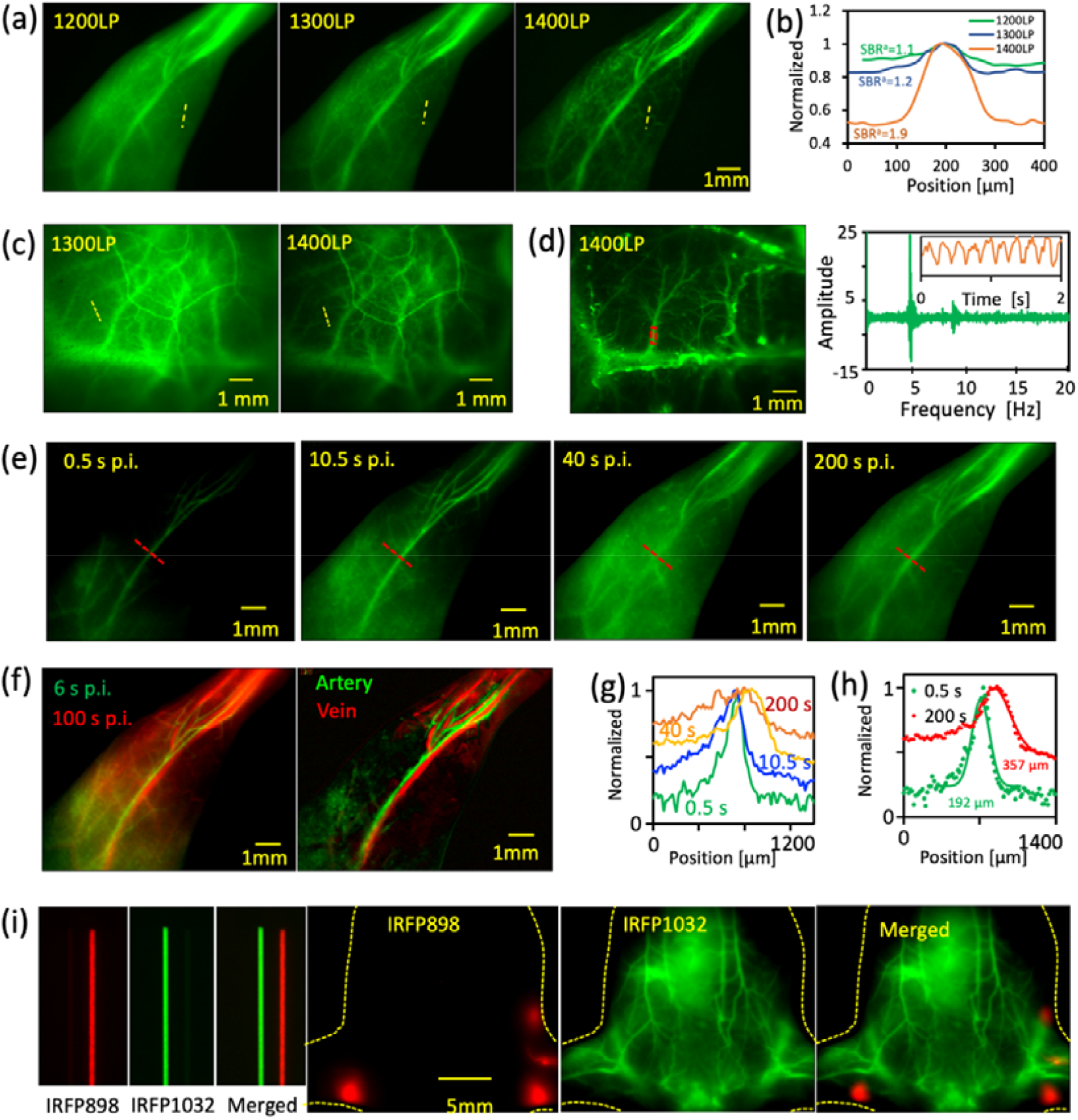
In vivo fluorescence imaging of IRFP1032 and IRFP898. (a) The in vivo fluorescence images of mouse hindlimb recorded with different long pass filters at 10 mins post intravenous injection of IRFP1032 liposome, excited at 980 nm. (b) The signal to adjacent background ratio (SBR^a^) of blood vessels of interest in (a) yellow dash line. (c) The through intact scalp and skull in vivo fluorescence imaging of mouse brain vessels recorded at 10 mins post intravenous injection of IRFP1032 liposome, excited at 980 nm, 75 mW/cm^2^ with 1300 nm and 1400 nm long pass filter, respectively. (d) The high temporal recording of blood flow fluctuation in brain vessel (location of vessels in red rectangle) with frame rate about 33 fps, exposure time 15 ms, and Fourier transform of the brain blood flow fluctuation in 50 seconds, with frequency about 4.3 Hz, heart beats about 258 beats per minute for the anesthetized mouse, inset: The blood flow fluctuation of brain vessel. (e) Real time imaging of blood flow dynamics with IRFP1032 liposome of hindlimb vessels at different time points p.i. (post injection) with a 1200 nm long pass filter, exposure time 50 ms, frame rate about 16 fps, excitation at 980 nm, 75 mW/cm^2^. (f) The artery and vein assignment at two time points 6 s p.i. and 100 s p.i. and the principal component analysis (PCA) of the real-time recorded fluorescence images. (g) The normalized fluorescence intensity of interest vessels shown in (e) red dash line. (h) The FWHM of main hindlimb artery (0.5 s p.i., saphenous artery) and vein (200 s p.i., great saphenous vein) by gaussian fit. (i) The dual color imaging of blood vessels and lymph nodes by labelling with IRFP1032 and IRFP898, respectively. All experimental details in supplementary information experimental section.

The excellent emission properties of IRFP1032 render the real time observation of the blood flow dynamics feasible. As shown in Figure 3d, the brain blood flow fluctuation induced by the heart beating could be evidently recorded and the corresponding heartbeat rate of the presented anesthetized mouse was found to be 258 beats per minute. Furthermore, the blood flow from the saphenous artery to capillary vessels, and then collected back through the great saphenous veins was unambiguously recorded via intravenous injection of the IRFP1032 liposome with frame rate about 16 fps (exposure time 50 ms) as shown in Figure 3e. Sooner post injection (p.i.), the saphenous artery could be clearly observed with very low background signal, with the signal to adjacent background (SBR^a^) is about 3.2 (Figure 3e 0.5 s p.i., Figure 3g green solid line, Figure S8a). As the blood flowing, the hindlimb muscle tissue was lit up by the perfusion of the IRFP1032 from the main vessels to capillaries (Figure 3e 10.5 s and 40 s p.i.). Around 40 s p.i., the saphenous artery was hardly distinguished from the surrounding tissue background, in spite of the fluorescence intensity at the location of the vessels not varying much during that time span (Figure S8b, Figure 3g orange solid line). In the end, the great saphenous veins (Figure 4a, 200 s p.i.) which is located next to the saphenous artery was gradually visualised (Video 1 in supplementary information). The relative location of arteries and veins could be obviously distinguished either by overlapping two images at different time points or by principal component analysis (Figure 3f, S9). The FWHM of the saphenous artery was around 192 μm which was much smaller than that of the great saphenous vein (357 μm, Figure 3h). Furthermore, the barely observed saphenous artery in late period p.i. (200 s p.i.) might be caused by the fact that the artery is located deeper than the vein, therefore the signal is covered by the scattering background (Figure S10).

**Figure 4.**
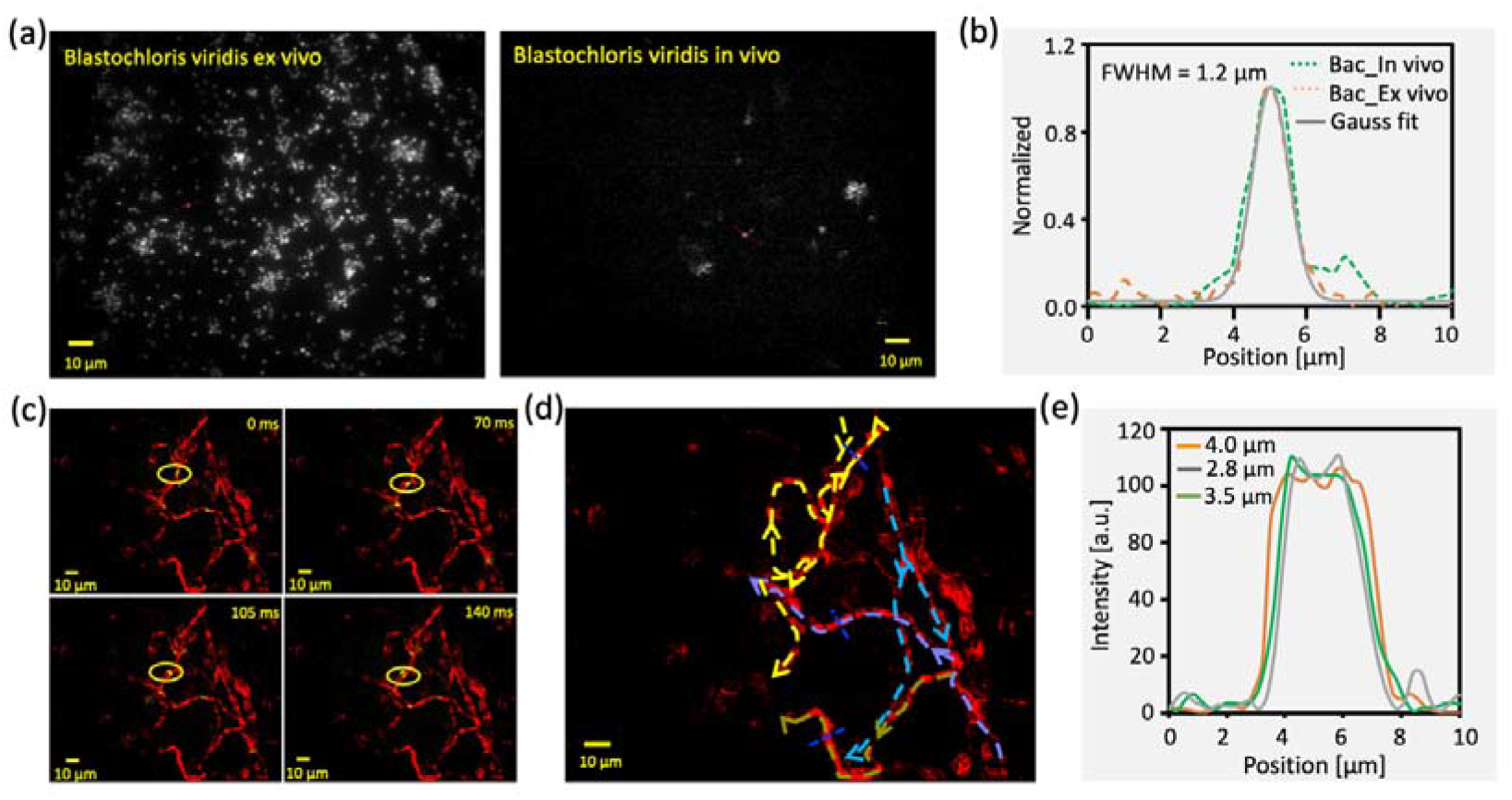
The single bacterial cell in vivo tracking. (a) The microscopy image of Blastochloris viridis, the Blastochloris viridis on glass slide (Left), the in vivo image of single bacterial cell in mouse ear blood vessel (Right), objective 60x, excited at 980 nm, laser intensity 75 mW/cm^2^, 1100 nm long pass filter, exposure time 25 ms. (b) The gaussian fit of the single bacterial cell as indicated by the red dash line in (a). (c) The motion of a cluster of bacterial cells in mouse ear blood vessels recorded at different times as shown in yellow circle, the red channel is constructed by projecting the maximum value of bacteria motion images in 120 seconds, green channel is the image of bacterial cell, microscopy setting as (a). (d) The blood flow direction of mouse ear vessels, the arrow indicates the flowing direction. (e) The cross section of the blood vessels of interest in (d) blue dotted lines. Experimental details in supplementary information experimental section.

As shown in Figure 3i, a dual color imaging with IRFP1032 and IRFP898 was performed to label the blood vessels and lymphatic network. The fluorescence maximum of IRFP898 is about 140 nm blue shifted to that of IRFP1032 (1032 nm, FWHM 70 nm) at around 898 nm with FWHM about 50 nm. Therefore, the well separated emission spectrum of both fluorescence proteins encouraged us to pursue a dual color in vivo imaging. Two well-known popliteal nodes could be distinctly observed at about 30 mins p.i. by intradermal injection of IRFP898 into the hind footpads.^45^ Surprisingly, a lymph vessel linking to the inguinal lymph node (ILN) and a blind-end lymph vessel were recorded on the left side of the mouse (lay on back). In general, the injection through the hind footpad would mainly drain to the popliteal lymph node (PLN), and a resection of PLN is necessary to visualize the ILN through hind footpad injection (Figure S11).^49^ The blood circulation system was labelled with IRFP1032, and the complex blood vessel network was clearly imaged with excitation at 980 nm and observation via 1200 nm long pass filter. The well resolved lymphatic and blood vessels network by labelling with IRFP898 and IRFP1032 respectively, manifested the outstanding potential for multiplex in vivo imaging with the fluorescence proteins.

### Single bacterial cell In vivo tracking

Taking the advantage of strong exciton absorption and emission of IRFP1032, the single Blastochloris viridis (Bv) cell could be tracked in real time. The Bv cell was intravenously injected after filtrating with porous membrane (pore size about 3 μm). A single Bv cell travelling through the capillaries of mouse ear was recorded as size about 1.2 μm which is similar to that on glass slide (Figure 6a, 6b, S12). By projecting the maximum value of each in vivo image (all over 3400 images taken within 120 s), the motion trajectory of the bacterial cell could be reconstructed, as shown in Figure 6c red channel. The trajectory diameter was found to be around 3.5 μm, which is in well agreement to the reported capillary size (Figure 6e).^50^ An interesting observation was tracked in real time that a cluster of Bv cells (probably about 3-5 bacterial cells, as shown in Figure S12f) was split into two parts while passing through a bifurcation of capillary shown in Figure 6c yellow circle. As indicated in the supplementary information Figure S13, a pair of Bv cells were tracked frame by frame with the relative distance remaining similar and no other bacterial cells jumped in among that time span. The tracked bacterial cell pair indicates that the motion direction of Bv cell in the presented anaesthesia mouse ear capillary could be unambiguously tracked at frame rate about 28.5 fps (Video 4 in supplementary information). Following the bacterial cell moving trajectory, very complex blood flow directions of mouse ear capillary were depicted as dash arrows in Figure 6d. For instance, the yellow arrow shows the blood first flowing forward then backward through the branch vessel.

### The expression of IRFP1032 in mamalian cells

The genetically encoded fluorescent protein as one of the most fascinating molecular tools, has been dramatically changing the approaches for biological research.^26,29^ However, currently developed genetically encoded fluorescence proteins are focused in the relatively strong tissue scattering region (visible and NIR-I). The excellent NIR-II exciton performance of the IRFP1032 discovered herein would shed light on the application of genetically encoded fluorescent protein for in vivo imaging. As we discussed above, the IRFP1032 is composed of *α, β, γ* polypeptide units and the polypeptide binding ligands (bacteriochlorophyll b and carotenoid).^39^ Moreover, the *α, β, γ* polypeptide units and the corresponding ligands of the IRFP1032 are controlled by different genes in wild Blastochloris viridis which makes the development of genetically encoded IRFP1032 a great challenge.^51^ In order to obtain the recombinant IRFP1032 in mammalian cells, the *α, β, γ* polypeptide units were expressed via a pBud plasmid and the binding ligands were delivered into cells via culture medium. To achieve a similar expression number of *α, β, γ* polypeptide in cells, a two promoter system and a self-cleaving peptide sequence were designed on a mammalian cell expression plasmid (pBud) as shown in Figure 5a. The CMV promoter was applied for the *γ* and *α* polypeptide, which were linked by a P2A self-cleaving peptide sequence. The EF1aphla promoter was chosen for the *β* polypeptide. The amino acids sequence of *α, β, γ* polypeptide were described in Figure 5b. While translating, the linked *γ* and *α* polypeptide might be broken off around the P2A linker.^52^ The IRFP1032 expression vector was first transient transfected to Hela cells via mixing with lipofectamine 2000, and then the polypeptide binding ligands in DMSO were added to the cell culture for incubation about 12 hours. The fluorescence images of the Hela cells with and without transiently transfected IRFP1032 vector were shown in Figure 5c. A fluorescence image was able to be observed among a few Hela cells in the group of expression vector transfected cells (excited at 980 nm and collected through 1000 nm long pass filter). While there is no sign of observable fluorescence for the vector (-) cells (without expression vector) under the same imaging condition and culture medium. The Hela cells were then centrifugal collected and imaged with 1000 nm long pass filter as shown in Figure 5d. A relative bright fluorescence image was able to be observed for the Hela cells with expression vector transfected (vector +). Furthermore, a differentiation could be observed between the vector (+) and vector (-) cells by subcutaneous injection into a nude mouse (fluorescence imaging under 1000 nm long pass filter, Figure 5e). In order to further confirm the fluorescence from the vector (+) cells was originated from the recombinant IRFP1032, the fluorescence spectrum was measured with excitation at 980 nm as shown in Figure 5f. Even though, the fluorescence intensity of the recombinant IRFP1032 in cells is relatively weak, the emission band of which was found to be in good agreement with the isolated IRFP1032 from Blastochloris viridis. The observed weak fluorescence from the Hela cells (expression vector transfected) indicates the poor expression of the IRFP1032. The low quantity of the recombinant IRFP1032 in Hela cells might be caused by the several parameters, for instance, the low expression efficiency of the *α, β, γ* polypeptide unit, the improper folding and assembling of the polypeptide helixes, etc. It might be necessary to optimize the amino acids sequence to reach a stable and high expression of IRFP1032 in mammalian cells.

**Figure 5.**
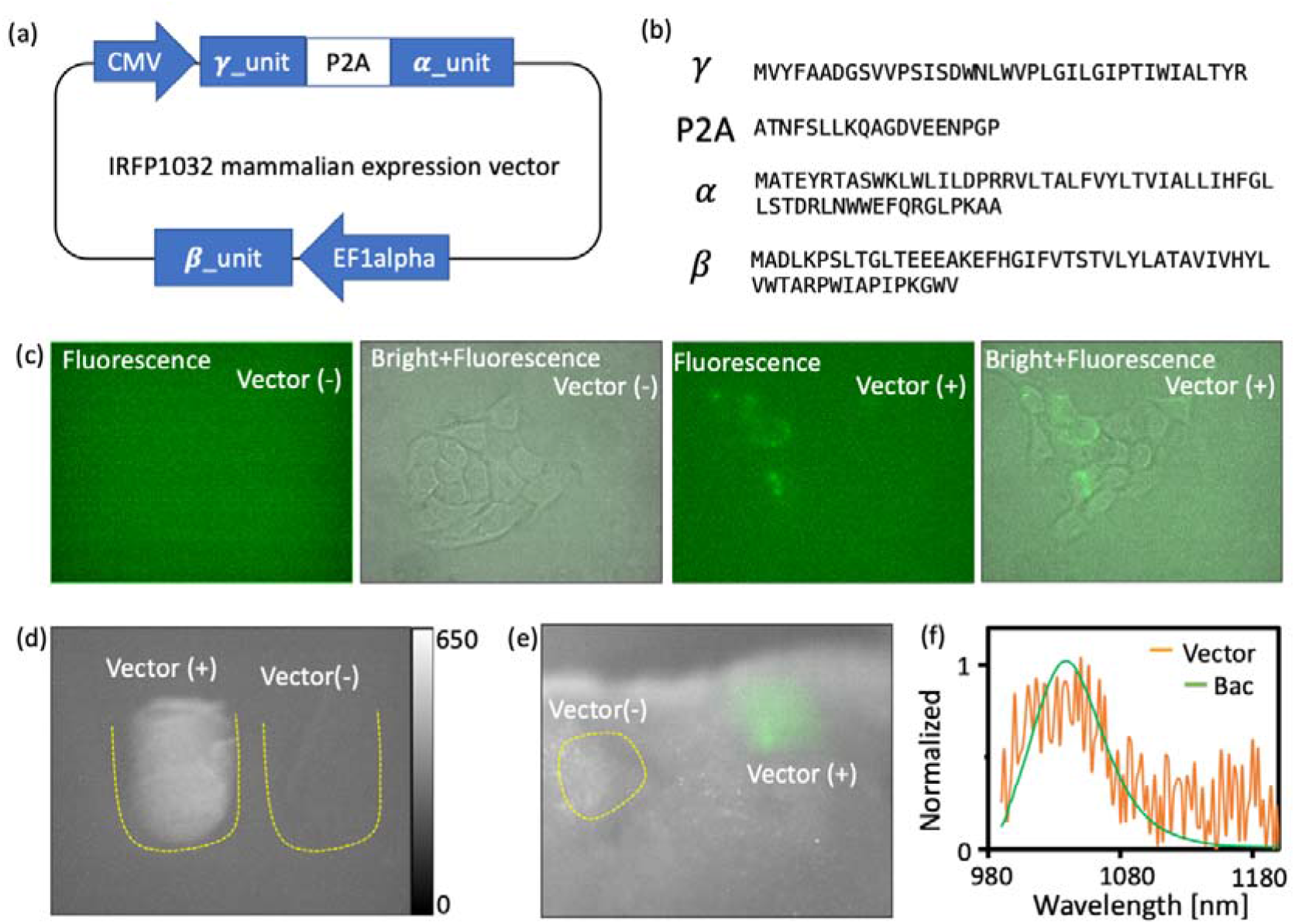
The expression of IRFP1032 in Hela cell. (a) The construction of the mammalian expression vector of IRFP1032 based on pBud plasmid. The CMV promoter for the *γ* and *α* polypeptide and the EF1alpha promoter for the *β* polypeptide of IRFP1032. (b) The amino acid sequence of the *α, β, γ* subunits of IRFP1032 and the P2A self-cleaving peptides sequence.^39^ (c) The image of the Hela cells with (Vector +) and without (Vector -) transfection of the IRFP1032 expression vector (cells incubated with ligands added culture medium, Excited at 980 nm, collected 1000 nm long pass filter, 60X objective, laser power 75 mW, exposure time 200 ms). (d) The image of the centrifugal collected Hela cells with (Vector +) and without (Vector -) transfection of the IRFP1032 expression vector (excited at 980 nm, collected 1000 nm long pass filter, laser power 25 mW, exposure 100 ms). (e) The fluorescence image of the subcutaneous injection of the centrifugal collected Hela cells with (vector +) and without (vector -) transfection of the IRFP1032 expression vector (excited at 980 nm, collected 1000 nm long pass filter, laser power 75 mW, exposure 200 ms). (f) The normalized emission spectra of the expressed IRFP1032 in Hela cell (orange curve) and IRFP1032 isolated from Blastochloris viridis (green curve).

### Conclusion and Outlooks

Here, we report the first fluorescent protein complex in the second near infrared window (IRFP1032). The excellent exciton absorption and emission properties of IRFP1032 render it to be the brightest NIR-II fluorophore ever reported (to the best of our knowledge). Taking advantage of the outstanding NIR-II performance of IRFP1032, high contrast through scalp and skull in vivo imaging was realized. Furthermore, the IRFP1032 also exhibits an excellent potential in real-time imaging, as illustrated by the video-rate speed recording of the brain blood flow fluctuation, the hemodynamic from artery to vein through capillaries, and single bacterial cell in vivo tracking. The IRFP898 as a second near infrared fluorescent protein with a well separated spectrum to IRFP1032 was also discovered. The blood and lymphatic networks were able to be clearly resolved by simultaneously applying both fluorescence proteins. Moreover, a mammalian expression vector was constructed for the IRFP1032 based on two promoter system. The NIR-II fluorescence from the recombinant IRFP1032 was able to be recorded unambiguously.

The promising NIR-II properties of IRPF1032 demonstrated here would open a new scene in fluorescence protein-based in vivo imaging. Since the discovery of fluorescence protein, the effort to pursue the longer wavelength emitting fluorescence proteins never has stopped. Transitioning to the near infrared fluorescence imaging, especially to the NIR-II region, is the most feasible way to realize high resolution and deep tissue in vivo imaging on biological studies, for instance the biological distribution of stem cells, neuron communication. Further work would be focused on optimizing the mammalian expression vector to reach a high and stable expression of IRFP1032. Furthermore, as the development of biosynthetic technology, the genome for the synthesis of chlorophyll has been successfully transferred and expressed in E. coli.^53^ To express the necessary synthetic enzymes of bacteriochlorophyll and carotenoids in the mammalian cell might be also possible, in which case the NIR-II fluorescence proteins of IRFP1032 might be served as convenient as the traditional fluorescence proteins.

## Methods

### Materials

All chemicals were used as received. Sodium succinate, K_2_HPO_4_, KH_2_PO_4_, MgSO_4_7H_2_O, (NH_4_)_2_SO_4_, CaCl_2_, 1,2-distearoyl-sn-glycero-3-phosphoethanolamine-N-[amino(polyethylene glycol)-2000](ammonium salt) (DSPE-PEG 2000), 1,2-dipalmitoyl-sn-glycero-3-phosphocholine (DPPC), Cholesterol, n-Dodecyl β-D-maltoside (DDM), Ferric Citrate, EDTA, chloral hydrate were purchased Aladdin chemistry. Yeast extract was purchased from Oxoid. Intralipid 20% was purchased from Sigma-Aldrich. IR26 was purchased from Exciton. Polycarbonate membrane with pore size 100 nm was purchased from Avastin. Polycarbonate membrane with pore size 3 μm was purchased from Millipore. Blastochloris viridis and Rhodospirillum rubrum were purchased from China General Microbiological Culture Collection Centre. The expression vector of IRFP1032 was constructed by GeneralBioL. lipofectamine 2000, culture mediums were purchased from ThermoFisher.

### Fluorescent protein preparation

The harvested bacteria were crushed through a probe sonicator (Amplitude 45%, 5s on and 5s off, vibracell) in a 40 ml PBS solution with adding MgCl_2_ (2.5mM, 1ml) under an ice bath for 30 mins. The cell membrane was first separated from the bacteria debris via centrifugation at 6000 rpm (Sigma 3k15) for 15 mins. The fluorescent protein isolation processes were modified from the reported procedures.^39^ The supernatant from the last step was further treated with ultra-centrifugation at force 110,000 g (Himac CR30NX) for 4 hours at 4 ^°^C. The fluorescent protein containing membrane was separated from the solution and collected by dissolving in PBS solution with adding MgCl_2_ (2.5mM, 1ml). The absorbance of the resulting solution was adjusted to about 20 at the exciton absorption maximum (1 cm optical path) and 3% (w/w) *n*-dodecyl β-d-maltoside (DDM) was added to the solution. The mixture was first gently mixed on a rocking mixer for 1.5 hours in the dark at room temperature and then further crushed through a probe sonicator (Amplitude 45%, 5s on and 5s off, vibracell) under an ice bath for 15 mins. The unsolublized components were removed from the mixture via centrifugation at 20,000g for 1 hour. Then about 1 ml of resulting supernate was added carefully on the top of a sucrose step gradient solution (from top to bottom 0.25 M, 0.5 M, 0.75 M, 1.0 M, 1.25 M in PBS buffer with 2% DDM). The sucrose solution was slowly added to the centrifugation tube along the wall by a long needle syringe from the high concentration to low concentration, each concentration with about 6 ml. After centrifugation at force 110,000 g (Himac CR30NX) for 16 hours at 4 ^°^C, the fluorescent protein was isolated as a coloured band and carefully collected via a syringe. The collected fluorescent protein was washed with a PBS buffer for 5 times under 100KD cut off centrifuge filter (Vivaspin) to remove the excess DDM at 6000 rpm (Sigma 3k15). Then the collected fluorescent protein was passed through a 300KD cut off centrifuge filter (Vivaspin). Finally, the isolated fluorescent protein was collected for further experiments.

### Expression of IRFP1032 in Hela cells

The IFRP1032 vector (500 ng) and Lipofectamine 2000 (2µL) were gently mixed with 50 µL of Opti-MEM, respectively, and incubated for 5 min at room temperature. Then the Opti-MEM solution of the IRFP1032 vector and the Lipofectamine 2000 were mixed and incubated for another 20 min at room temperature. The Hela cells with 70-90% confluency was divided into two groups, vector (+) and vector (-). For the vector (+) group, 100 µL of vector-Lipofectamine 2000 mixed Opti-MEM was added. For the vector (-) group, 100 µL of fresh Opti-MEM was added. After 4 hours of incubation, the culture medium was replaced with a fresh complete cell culture medium and 100 µL ligands in the DMSO solution were added to both groups. Ligands were directly extracted by DMSO from isolated IRFP1032 with absorbance at 795 nm about 10. After another 12 hours incubation, the cell culture medium was replaced by PBS. The cells are then used for imaging, centrifugal collection, and subcutaneous injection.

### Photo-properties characterization

The NIR fluorescence spectrum was taken with Edinburgh Instruments FLS920. A CW diode laser was used to pass through the solution sample in quartz cuvette with optical path 1 cm. The emission was collected by an InGaAs detector from 850 nm to 1500 nm with detector response correction. The absorption spectra were recorded by UV-2700 and UV-3600 (Shimadzu) with quartz cuvette (optical path 1 cm). See Supplementary Information for the calculation of the quantum yield and extinction coefficient.

### NIR-II in vivo imaging setup

A home-built cooling system was used to deep cooling an InGaAs camera (650 × 512 pixels, Tekwin, designed with cooling temperature −10 ^°^C) to −30 ^°^C, the gain was set to high, and different exposure times were used to achieve sufficient signal and frame rates. A 1 W, 825 nm diode laser and a 5 W 980 nm diode laser were used to produce the excitation beam, which was coupled into a core size 400 μm metal cladded multimode fiber (Thorlabs), collimated by a collimator (980 nm, f = 6.29 mm, NA = 0.37, Fiber Collimation, Thorlabs) and then diffused through an engineered diffuser (50° Circle Pattern Diffuser, Thorlabs) for generating a homogeneous illumination, the power intensity for 825 nm about 25 mW/cm^2^ and for 980 nm about 25 or 75 mW/cm^2^ were used. A single lens system was used to focus the image on camera (Achromatic Doublet, 1050 - 1700 nm, focal length 50 mm, Thorlabs), the lens position was adjusted to achieve a different field of view. Short pass filter was used on the excitation beam path and various long-pass and short pass filters were incorporated into the collection path. Fiji was used for all image treatments.

### NIR-II epi fluorescence microscopy

A home-built microscopy system with the above mentioned InGaAs camera was used for microscopic imaging. A diode laser (5 W, 980 nm) was used to produce the excitation beam, which was coupled into a core size 400 μm metal cladded multimode fiber (Thorlabs), collimated by a collimator (980 nm, f = 6.29 mm, NA = 0.37, Fiber Collimation, Thorlabs), then diverged through an achromatic doublet lens (focal length 150 mm, Thorlabs) before reaching the objective (60x, APO, NIR, Nikon), power intensity about 75 mW/cm^2^ were used. A long pass dichroic mirror (cut-on 1000 nm, Thorlabs) was used for collimating the excitation beam and fluorescence. An infinity-Corrected tube lens with focus length about 200 mm (Broadband MgF_2_ Coating, Thorlabs) was used to focus the image on the camera. Short pass filters were used on the excitation beam path and various long pass filters were incorporated into the collection path. Fiji was used for all image treatments (maximum intensity projection, pixel intensity, etc.).

## Acknowledgements

We would like to thank Dr. Jun Chu and Dr. Feng Liu for the help of design the expression vector. This work was supported by the National Natural Science Foundation of China (No. 22077135, No. 21905296).

## Author contributions

Qinchao Sun conceived and designed the experiments. Qinchao Sun, Zong Chang, ChenChen Liu, Shubi Zhao, Jiaqi, Chen, Hanyu Tian performed the experiments. Qinchao Sun and Zong Chang analyzed the data and wrote the manuscript. All authors discussed the results and commented on the manuscript.

## Additional information

Supplementary information is available in the online version of the paper. Correspondence and requests for materials should be addressed to

