## supplemental file for "Near Infrared-II Fluorescent protein for In-vivo Imaging"

**Experimental Section**

**Bacteria culture.** Blastochloris viridis and Rhodospirillum rubrum were cultured with the same medium (ATCC 19567). Briefly, a culture medium of sodium succinate (2.5 g), K_2_HPO_4_ (0.9g), KH_2_PO_4_ (0.6 g), MgSO_4_ 7H_2_O (0.2 g), (NH_4_)_2_SO_4_ (1.25 g), CaCl_2_ (0.07 g), ferric citrate (3.0 mg), EDTA (2.0 mg), yeast extract (0.5 g) and distilled water 1.0 L were autoclaved at 120 ^o^C for 15 minutes, and then 3% of cysteine was added after cooling to room temperature. 20 μl of the corresponding bacteria storage was added and the bottle was sealed to incubate at 30 ^o^C under 100 W Tungsten Halogen lamp. The bacteria were harvested after 5 – 10 days of incubation.

**Liposome preparation.** A solution of IRFP1032 with OD about 30 at 980 nm (1 cm optical path) was mixed (1:1 in volume) with DPPC and DSPE-PEG2000 PBS solution which was first dissolved in 1 ml CHCl_3_ (DPPC 50 mg, DSPE-PEG2000 5 mg) then dried off under vacuum and resuspended in 4 ml PBS under sonication bath. The mixture was passed through a 100 nm membrane pore for a few times with the Avestin liposome setup. The final liposome solution of IRFP1032 with absorbance about 15 at 980 nm (1 cm optical path). Similar procedure was done for the preparation of IRFP898. A solution of IRFP898 with OD about 5 at 825 nm (1 cm optical path) was mixed (1:1 in volume) with DPPC and DSPE-PEG2000 PBS solution which was first dissolved in 1 ml CHCl_3_ (DPPC 50 mg, DSPE-PEG2000 5 mg) then dried off under vacuum and resuspended in 4 ml PBS under sonication bath. The mixture was passed through a 100 nm membrane pore for a few times with the Avestin liposome setup. The final liposome solution of IRFP898 with absorbance about 2.5 at 825 nm (1 cm optical path). The hydrodynamic size of the liposome was about 140 nm by DLS. All the imaging experiments were conducted with the liposome-based fluorescent protein. The emission and absorption experiments were done with the liposome-based fluorescent protein.

**Absorption and emission spectra.** The absorption spectra of bacteria chlorophyll a and b in methanol, IRFP898 in PBS and IRFP1032 in PBS were measured by UV-2700 and UV-3600 in a quartz cuvette with optical path 1 cm. The emission spectra of IRFP898 in PBS were measured by a calibrated InGaAs detector (Edinburgh Instruments FLS920) with absorbance at 825 nm lower than 0.1, optical path 1 cm, excited by diode laser with emission wavelength around 825 nm, power intensity about 25 mW/cm^2^. The emission spectra of IRFP1032 in PBS were measured by a calibrated InGaAs detector (Edinburgh Instruments FLS920) with absorbance at 980 nm lower than 0.1, optical path 1 cm, excited by diode laser with emission wavelength around 980 nm, power intensity about 10 mW/cm^2^.

**Relative fluorescence quantum yield.** The IR26 in DCM solution was used as reference for IRFP1032, IR820 in EtOH solution was used as reference for IRFP898. Five solutions with different absorbance (lower than 0.1) were prepared for reference dye and the protein fluorophores in PBS, respectively. For example, the absorbance at 980 nm was recorded for all the solutions in quartz cuvette (optical path 1 cm). The emission spectra were recorded from 1000 nm to 1500 nm by excitation at 980 nm with power intensity about 10 mW/cm^2^. The integrated area of IRFP1032 and IR26 from 1000 nm to 1500 nm was plotted as a function of the absorbance. A linear fitting was applied to both IRFP1032 and IR26. The relative quantum yield of IRFP1032 was calculated in following equation:

$$\frac{{QY}_{IRFP1032}}{{QY}_{IR26}}=\frac{n_{H2O}^{2}}{n_{DCM}^{2}}\times\frac{{Slope}_{IFRP1032}}{{Slope}_{IR26}}$$

The refractive index of DCM is taken as 1.4 and that of H_2_O is taken as 1.3. The slope is from the linear fitting function of corresponding fluorophores, IRFP1032 in PBS and IR26 in DCM. Same procedure was applied for the measurement of IRFP898. The Errors were calculated with three groups of QY measurements, for example, fifteen solutions of fluorophore prepared and divided into three groups to get the QY.

**Extinction coefficient measurements:** The key point of the measurement of extinction coefficient of IRFP1032 and IRFP898 was to get the accurate molar concentration of the fluorescent protein. As the surfactant and buffers used for purification of the membrane protein, the concentration of the IRFP1032 and IRFP898 was calculated by the concentration of bacteriochlorophylls. 100 μL of stock solution of fluorescent protein was first dried under N_2_ and then 1 mL of DMSO was added to extract the bacteriochlorophylls. The absorbance of the extracted bacteriochlorophyll was measured, and the concentration of the bacteriochlorophylls of the stock fluorescent protein was calculated by the extinction coefficient of bacteriochlorophyll. The concentration of corresponding fluorescent protein was calculated by the fact that 32 bacteriochlorophyll a in one IRFP898 molecule and 34 bacteriochlorophyll b in one IRFP1032 molecule. The extinction coefficient of bacteriochlorophylls was measured with extraction and purification of the bacteriochlorophylls via thin-layer chromatography. The measured extinction coefficient of bacteriochlorophylls was in good agreement as reported.^1,2^

**Animal handling.** All animal experiments were conducted in accordance with National Institutes of Health guidelines, and procedures were approved by the Animal Committee. Eight-week-old male BALB/c nude mice were obtained from Beijing Vital River Laboratory Animal Technology Co., Ltd. Before imaging, all mice were anaesthetized by intraperitoneal injection of chloral hydrate (400 mg/kg), and maintained in sufficient depth of anaesthesia, reinjection with one-third of the original dose as necessary. For Tail vein injection of contrast agents was carried out with an injection dose of about 150 μL in the PBS solution at specified concentrations. No blinding or randomization was required for the animal studies.

**Tissue Phantom Imaging.** Tissue phantom was prepared by 2.5% Intralipid (diluted from 20 % Intralipid) solution. Images were taken by covering a capillary of IRFP1032 in one well of a 12 well plate with 0.5 ml the prepared tissue phantom (about 1.4 mm depth, well diameter about 2.2 cm). For the 1100 nm long pass filter experiment, camera exposure time is 100 ms, excitation at 980 nm (5 mW/cm^2^). For the 1200 nm long pass filter experiment, camera exposure time is 100 ms, excitation at 980 nm (10 mW/cm^2^). For the 1300 nm long pass filter experiment, camera exposure time is 500 ms, excitation at 980 nm (75 mW/cm^2^). For the 1400 nm long pass filter experiment, camera exposure time is 1000 ms, excitation at 980 nm (75 mW/cm^2^). The imaging setup as described in the main text.

**In vivo imaging.** The Eight-week-old male BALB/c nude mouse was anesthetized, and 150 μL of IRFP1032 (through 0.22 μm syringe filter) with OD at 980 nm about 10 (optical path 1 cm) was injected through the tail vein. For real time blood flow dynamic imaging, the camera was set before injection, exposure time 50 ms, excitation at 980 nm (75 mW/cm^2^), collected through a 1200 nm long pass filter (Figure 3e). For brain blood fluctuation imaging, scalp removed mice were used and camera exposure time 15 ms, excitation at 980 nm (75 mW/cm^2^), collected through a 1200 nm long pass filter (Figure 3d). For brain blood vessel imaging, intact scalp and scalp removed mice were used and camera exposure time 1000 ms, excitation at 980 nm (75 mW/cm^2^), collected through 1300 nm and 1400 nm long pass filter (Figure 3c). For hind limb vessel imaging, camera exposure time is 1000 ms, excitation at 980 nm (75 mW/cm^2^) for collecting through 1300 nm and 1400 nm long pass filter, and camera exposure time is 200 ms, excitation at 980 nm (25 mW/cm^2^) for collecting through 1200 nm long pass filter (Figure 3a). Imaging setup as described in the main text.

**Dual color imaging.** The Eight-week-old male BALB/c nude mouse was anesthetized, and then 10 μL of IRFP898 (through 0.22 μm syringe filter) with OD at 825 nm about 1 (optical path 1 cm) was injected s.c. in the left and right hind footpad of the mouse. After 30 mins, 150 μL of IRFP1032 (through 0.22 μm syringe filter) with OD at 980 nm about 10 (optical path 1 cm) was injected through the tail vein. The image was taken sooner after the second injection, for imaging lymph vessels and nodes, excited at 825 nm (25 mW/cm^2^), collected through 900 nm long pass filter and 1000 nm short pass filter, camera exposure time 100 ms; for imaging blood vessels, excited at 980 nm (25 mW/cm^2^), collected through 1200 nm long pass filter, camera exposure time 100 ms. Imaging setup as described in the main text.

**Single bacteria imaging.** The Blastochloris viridis in PBS solution was passed through a 3 μm syringe filter before injecting (100 μL, absorbance at 980 nm about 1) via the tail vein into the anesthetized eight-week-old male BALB/c nude mouse, and the fluorescence was immediately imaged. The mouse ear was placed under the objective (60x, NIR, Nikon) of the home-built microscopy as described in the main text, excited at 980 nm (75 mW/cm^2^), collected through 1100 nm long pass filter, camera exposure time 25 ms.

**Image dimension calibration.** For a single lens imaging system, a ruler was imaged by Tungsten Halogen lamp illumination. For the microscopy system, the USAF resolution target was imaged by the transmission light from a Tungsten Halogen lamp.

**Principal component analysis.** PCA was performed by the built-in PCA function of MATLAB on a time stack of images for which pixel intensity varied as a function of time. The output coefficient was further separated by sign, as the positive coefficient and the negative coefficient. The distinguishing of arteries and veins was based on the fact that the fluorescence in the artery vessels of hindlimb should be observed first, since the intravenous injected fluorophores was first carried back to the lung and then pumped to the arteries by heart.

**Microscopy imaging through cranial window.** The implantation of the cranial window was similar as previously reported.^3^ Briefly, the head of the anesthetized mouse was fixed by a stereotactic apparatus. The head hair was removed and then the skin was cleaned with 75% ethanol solution. The skin was removed, and a 6-mm circle was drawn over the parietal regions of the skull, with a high-speed air-turbine drill. A groove circle was carefully made until the bone flap became loose. During the drilling process, cold saline was used to avoid thermal injury. After removal of the bone, an 8-mm cover glass was used to seal the window by glass ionomer cement. The prepared cranial window was placed under the objective (20x, Nikon) of the home-built microscopy as described in the main text, excited at 980 nm (25 mW/cm^2^), collected through 1200 nm long pass filter, camera exposure time 100 ms.

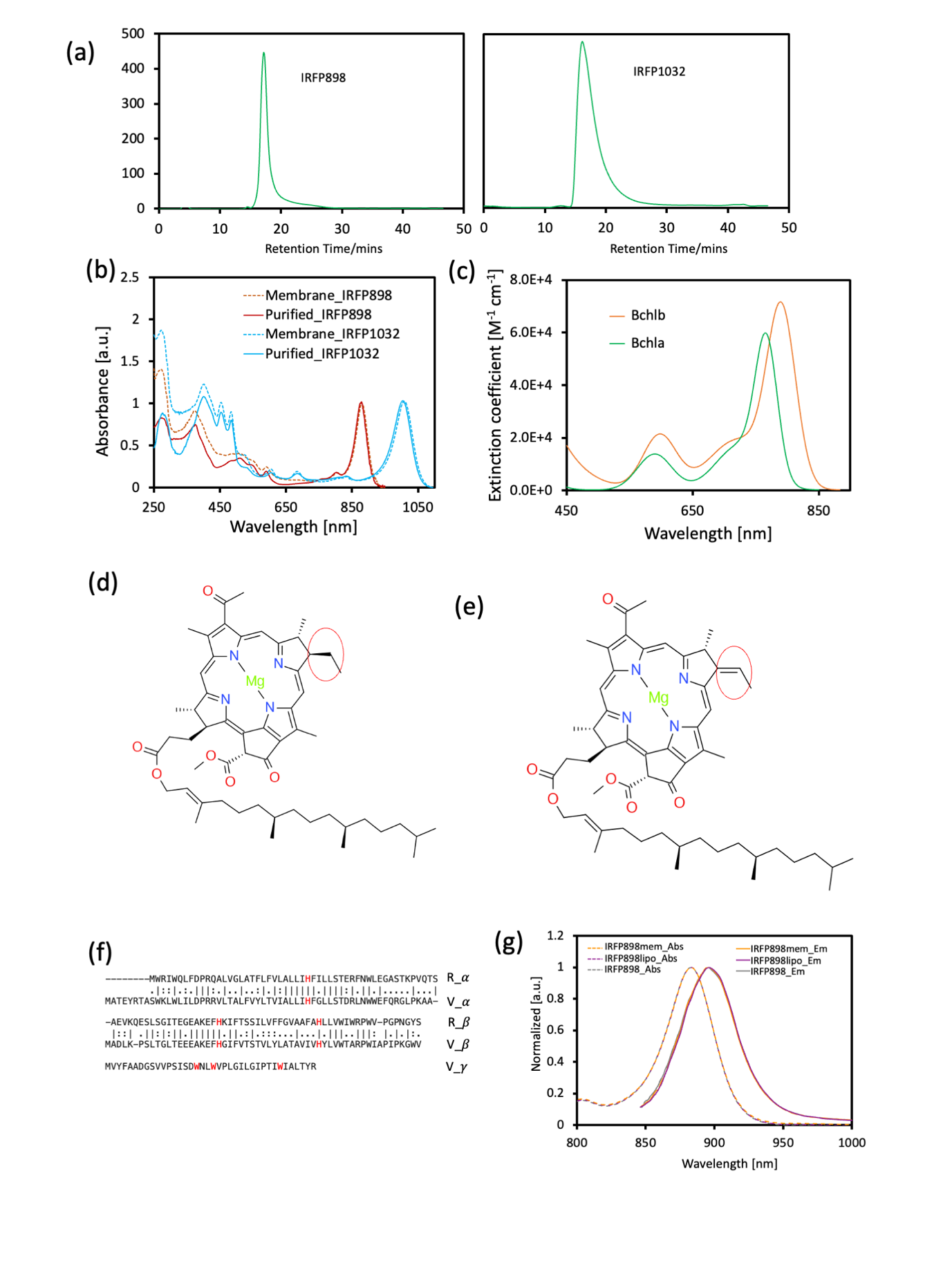

**Figure S1.** (a) The retention time of IRFP1032 and IRFP898 by HPLC. (b) The absorption spectra of fluorescent protein before and after purification from 250 nm to 1100 nm. The blue solid curve is the absorption of purified IRFP1032, the blue dash curve is the absorption of membrane IRFP1032, the red solid curve is the absorption of purified IRFP898 and the red dash curve is the absorption of membrane IRFP898. (c) The extinction coefficient of bacteriochlorophyll a and bacteriochlorophyll b in DMSO solution. (d) The molecular structure of bacteriochlorophyll a. (e) The molecular structure of bacteriochlorophyll b. (f) The amino acid sequence of the 𝛼, 𝛽, 𝛾 subunits of light harvesting protein complex of Blastochloris viridis (V_𝛼, V_𝛽, V_𝛾), the 𝛼, 𝛽 subunits of light harvesting protein complex of Rhodospirillum rubrum (R_𝛼, R_𝛽), and the corresponding alignment to the histidine (red). The vertical slash represents identical amino acid, the colon represents the similar properties of amino acid, and the dot represents less similarity. (g) The normalized absorption and emission spectra of IRFP898 protein in different environments, the IRFP898mem as in cell membrane (before isolation), IRFP898 as isolated from membrane by DDM, the IRFP898lipo as in liposome.

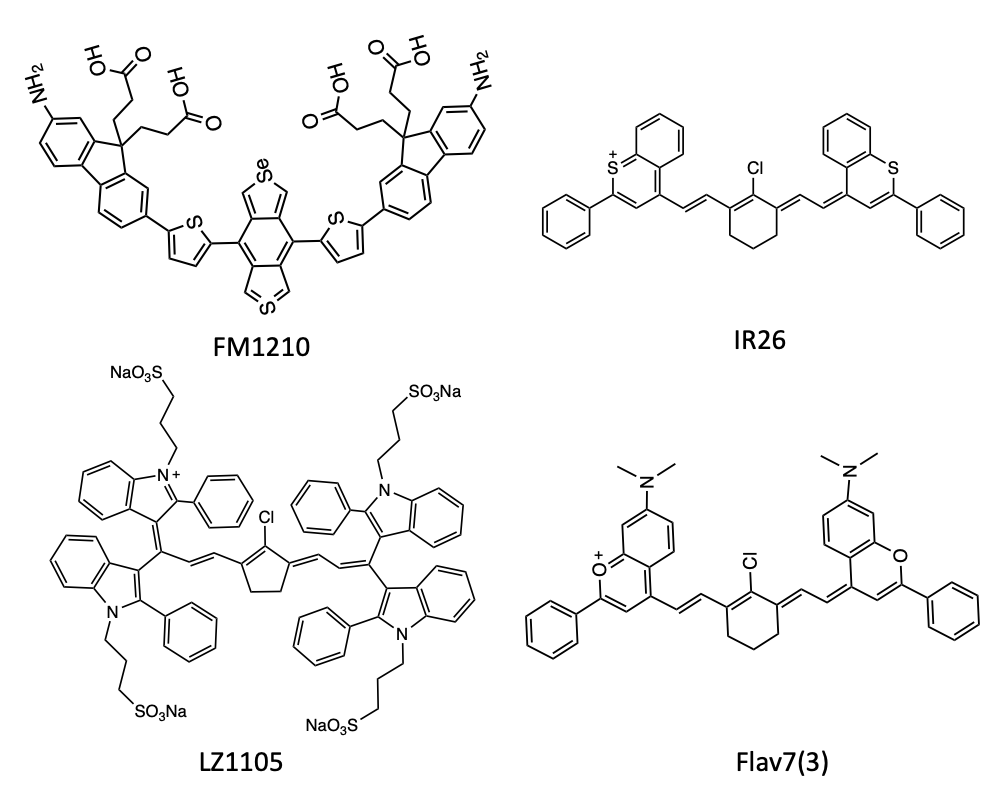

**Figure S2.** The molecular structure of several reported organic dyes with absorption and emission maximum in NIR-II or close to NIR-II region.

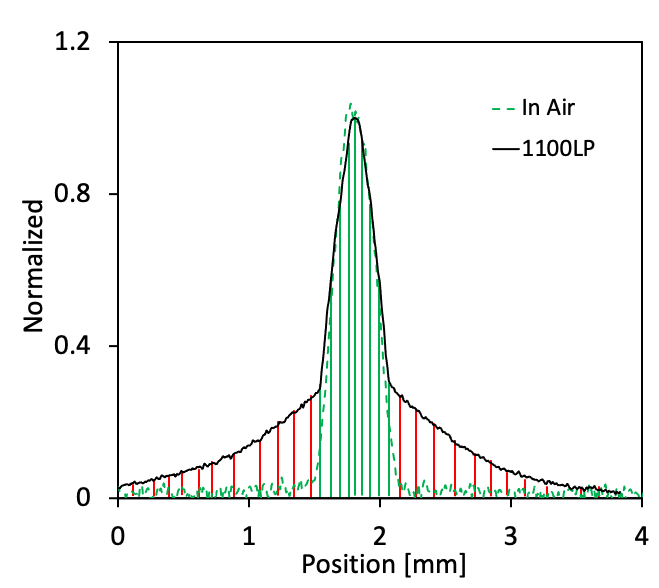

**Figure S3.** The black solid curve is the scattering pattern of the capillary (the fluorescence intensity as a function of position on cross section of capillary, Figure 3b yellow dash line) filled with IRFP1032 covered by 1.4 mm Intralipid phantom (2.5%). The green dotted curve is the fluorescence intensity as a function of position on the cross section of capillary filled with IRFP1032 in air (no intralipid phantom).

**The calculation of the ratio of scattered light by passing through 1.4 mm Intralipid phantom (2.5%).**

The non-scattered fluorescence photons emitted from a capillary in air forms the pattern as shown in Figure S3 the dotted green curve. Therefore, we consider the green solid line region as the contribution from the non-scattered photon or the photons undergoing a few scatterings but still effective for imaging in the case that the capillary is covered by 1.4 mm Intralipid phantom (2.5%). On the other hand, the red line region is considered to be contributed by the light undergoing dramatic scattering which causes the background. An integration was taken for both green line region (non-scattered) and red line region (scattered), respectively. The scattered photon portion could be reasonably calculated as follows.

R = A_R_/A_T_

Where A_R_ is the area of the red line region, A_T_ is the sum of the area of the red line region and the green line region.

**Table S1.** The scattered photon ratio measured at different long pass filters of the capillary filled with IRFP1032 covered by 1.4 mm Intralipid phantom 2.5%. We should notice that the ratio of scattered photon at the same wavelength highly depends on the scattering medium.

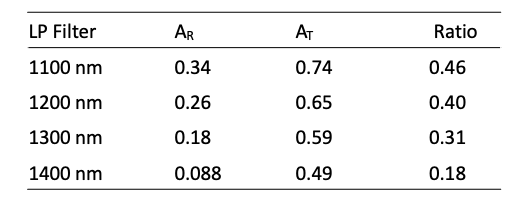

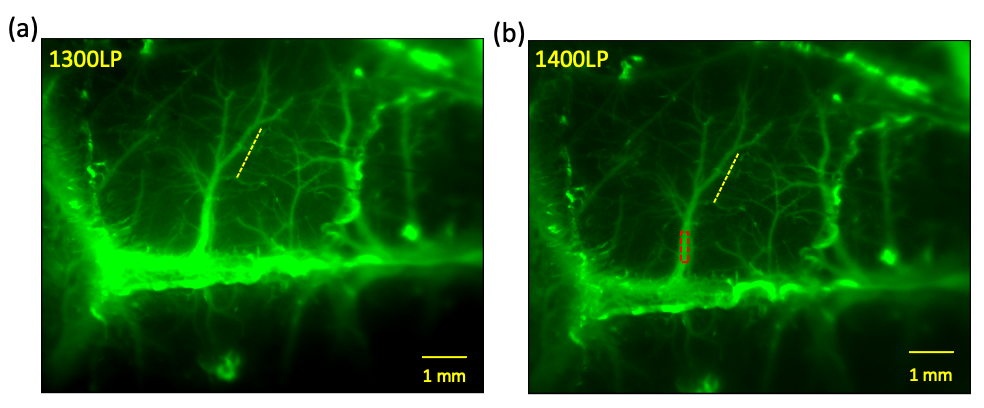

**Figure S4.** (a) The through skull (removed scalp) in vivo fluorescence imaging of mouse brain vessel recorded with 1300 nm long pass filter, intravenous injection of IRFP1032 liposome, excited at 980 nm, laser power 75 mW/cm^2^, exposure time 1000 ms. (b) The through skull (removed scalp) in vivo fluorescence imaging of mouse brain vessel recorded with 1400 nm long pass filter, intravenous injection of IRFP1032 liposome, excited at 980 nm, laser power 75 mW/cm^2^, 1000 ms.

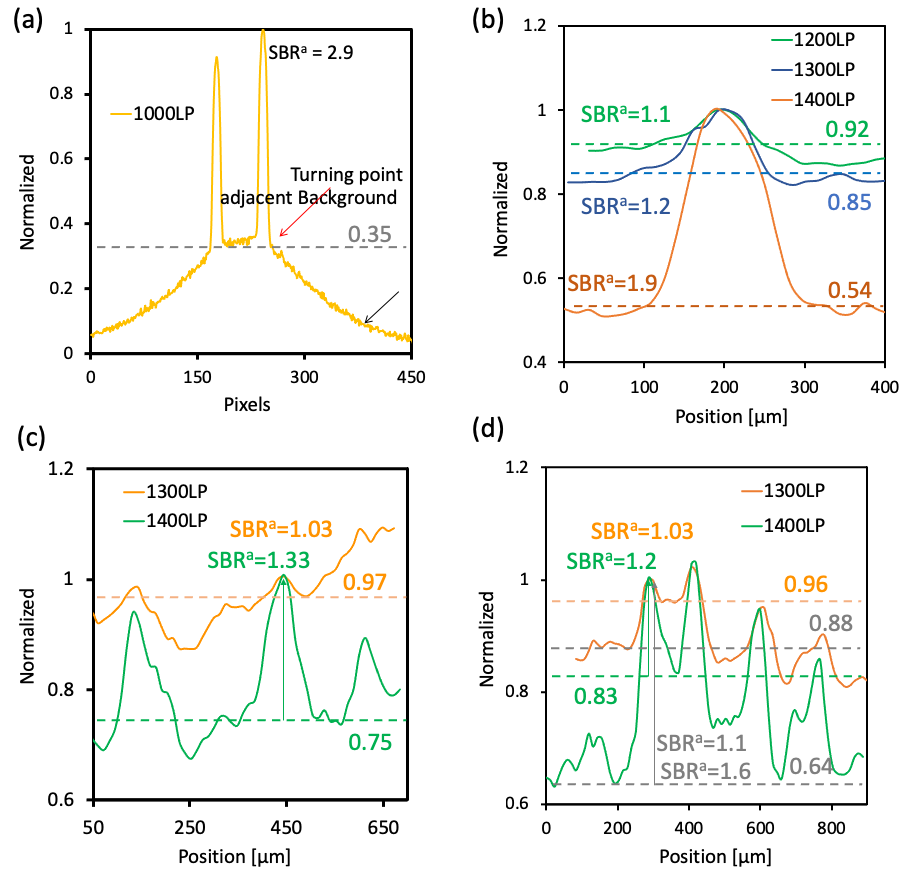

**Figure S5.** The signal to adjacent background ratio (SBR^a^). (a) The fluorescence intensity as function of pixels of two capillaries filled with IRFP1032 with slightly different concentration to mimic the two closely located vessels, excited at 980 nm, covered by 1.4 mm Intralipid phantom (2.5%), measured with 1000 nm long pass filter. (b) The SBR^a^ of the blood vessels as indicated by the yellow dash line in Figure 3d, the dash line indicates the turning point value of the vessel of interest. (c) The SBR^a^ of the blood vessels as indicated by the yellow dash line in Figure 3f, the dash line indicates the turning point value of vessels of interest, the arrow indicates the vessels of interest. (d) SBR^a^ of the blood vessels as indicated by the yellow dash line in Figure S4a, b, the arrow indicated the vessels of interest, the orange and green dash line indicated the higher turning point value of vessels of interest, the grey dash line indicated the lower turning point value of vessels of interest.

**The calculation of the signal to adjacent background ratio (SBR^a^).**

The calculation of the signal to background ratio of an image is kind of tricky, as the background could be set to close to zero, in which case the signal to background ratio could be huge and varies on image processing procedure by the researchers. Therefore, the signal to background calculation in present work, was done only by the subtraction of the dark counts of the camera, no further scaling and background setting were taken. As demonstrated in Figure S5a, the narrow band peaks correspond to the non-scattered photon imaging, and the broad band at the bottom comes from the scattered photons. It is reasonable to define the signal to background ratio as the ratio of the peak intensity divided by the value at the turning point as indicated by the red arrow (adjacent background), rather than set the background to a faraway point as indicated by the black arrow. In present work, the signal to background ratio was calculated by setting background at the turning point of the interested signal.

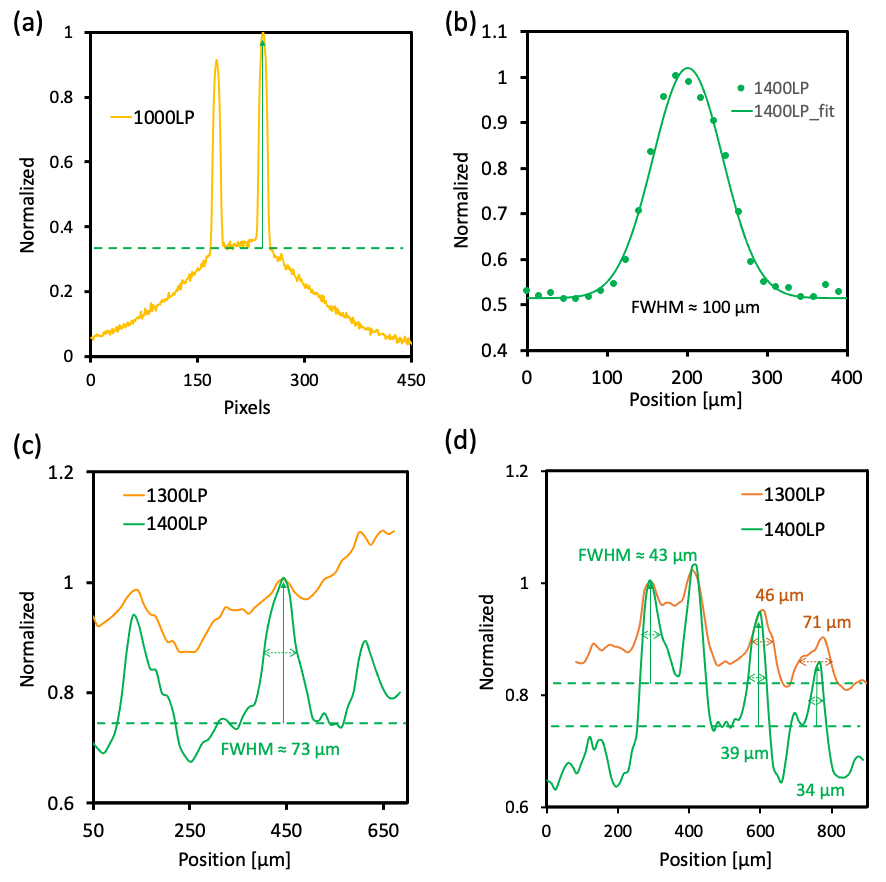

**Figure S6.** The FWHM of the vessels in the region of interest. (a) The fluorescence intensity as function of pixels of two capillaries filled with IRFP1032 with slightly different concentration to mimic the two closely located vessels, excited at 980 nm, covered by 1.4 mm Intralipid phantom (2.5%), measured with 1000 nm long pass filter. (b) The FWHM of a vessel of interest as indicated in the yellow dash line in Figure 3d(1400LP) by gaussian fit. (c) The FWHM of vessels of interest as indicated in the yellow dash line in Figure 3f. (d) The FWHM of vessels of interest as indicated in the yellow dash line in Figure S4a,b.

**The calculation of the FWHM.**

As a matter of fact, the high scattering effect and closely located vessels, in most of the case, the shape of the recorded vessels is not well gaussian shaped, as shown in Figure S6a. Therefore, it is reasonable to take the half maximum from the peak maximum to the turning point, as indicated by the green dash line in Figure S6a. For the vessels with well resolved shape and well separated from the other vessels, were fitted by gaussian function with FWHM =2.36𝝈

$${y=B+Ae}^{(-\frac{x-x_{0}}{2\sigma^{2}})}$$

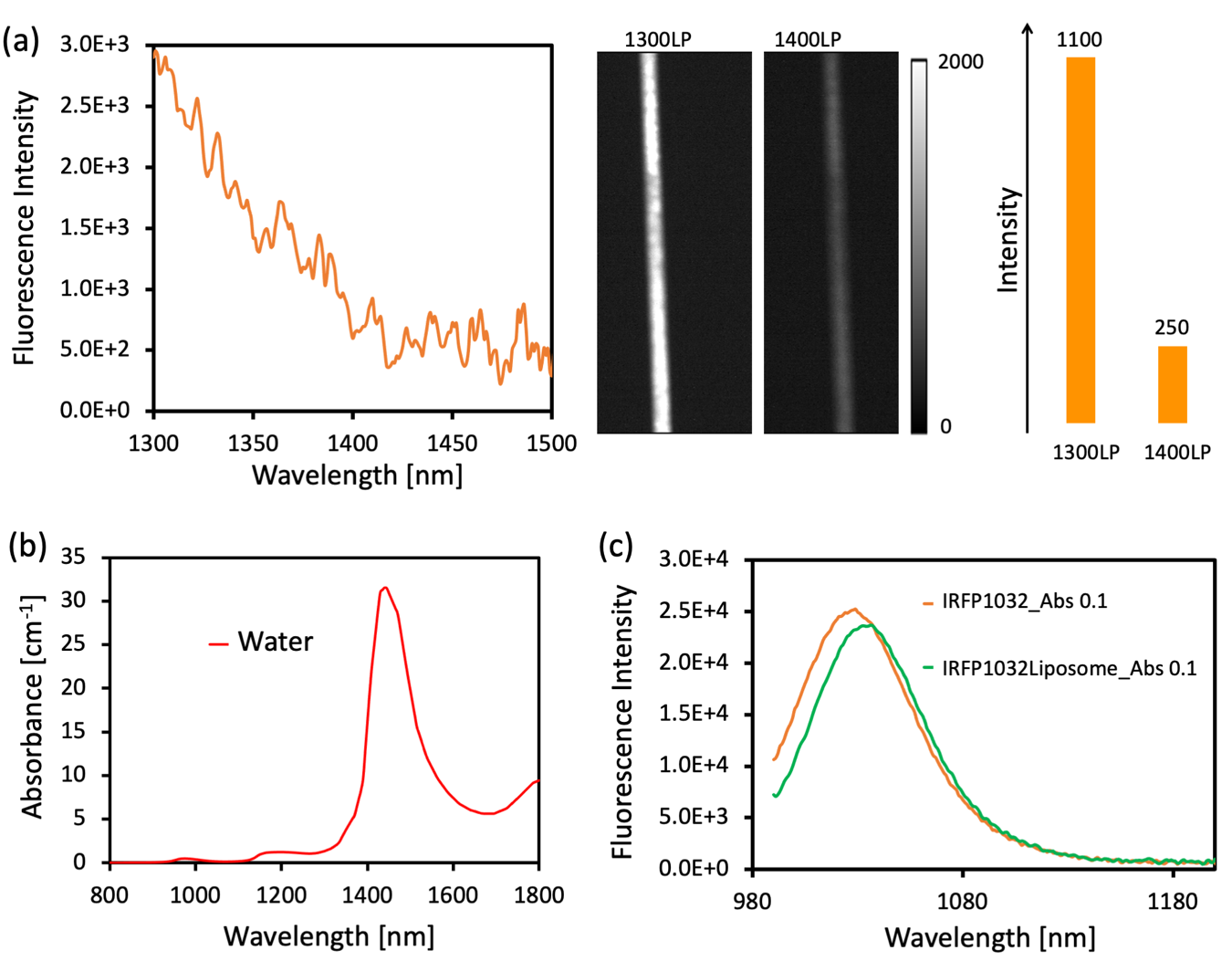

**Figure S7.** (a) The emission spectrum and intensity of IRFP1032 from 1300 nm to over 1400 nm. (b) The absorbance of water with an optical path 1 cm from 800 nm to 1800 nm. (c) The emission spectra of IRFP1032 with and without liposome treatment at absorbance at 980 nm about 0.1.

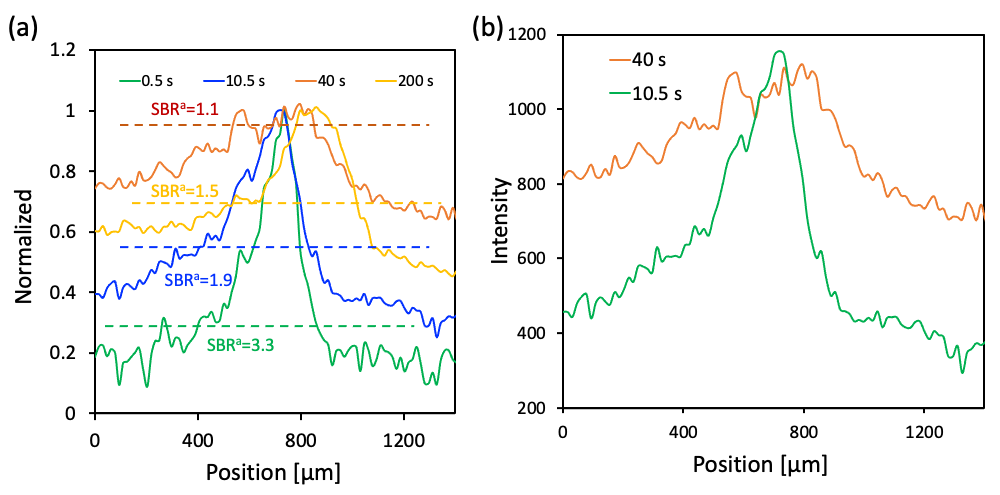

**Figure S8.** (a) The SBR^a^ at different times post injection of the vessels of interest, as shown by the yellow dash line in Figure 4a. (b) The fluorescence intensity of the vessels of interest as indicated by the yellow dash line in Figure 4a at 10.5 s and 40 s post injection.

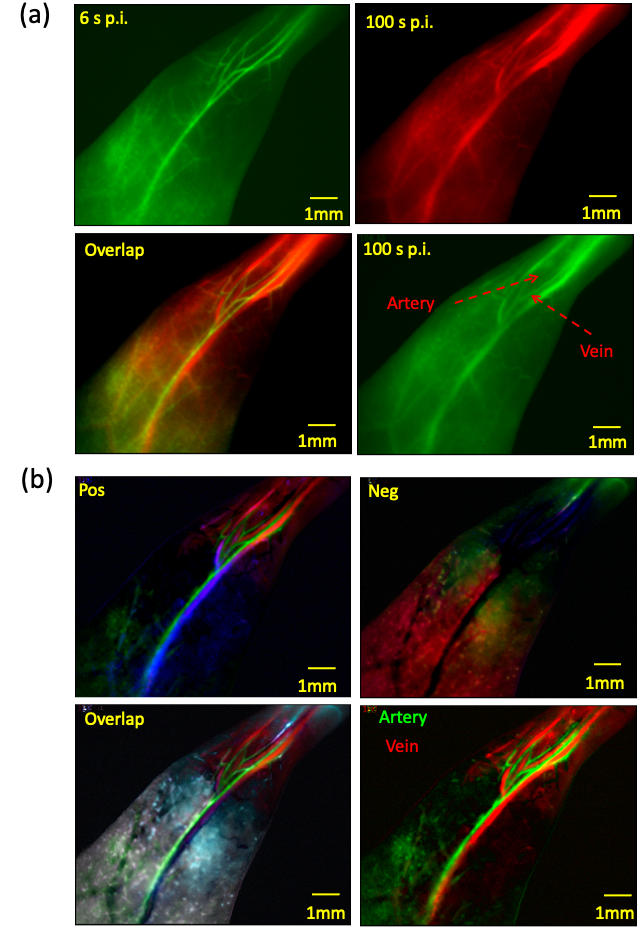

**Figure S9.** (a) The artery and vein assignment at two time points 6 s p.i. in green channel and 100 s p.i. in red channel. (b) The principal component analysis (PCA) of the real-time recorded fluorescence images. Three components were chosen for both positive (in RGB) and negative (in RGB), the artery was assigned with the first and third positive component, the vein was assigned with second positive and third negative component.

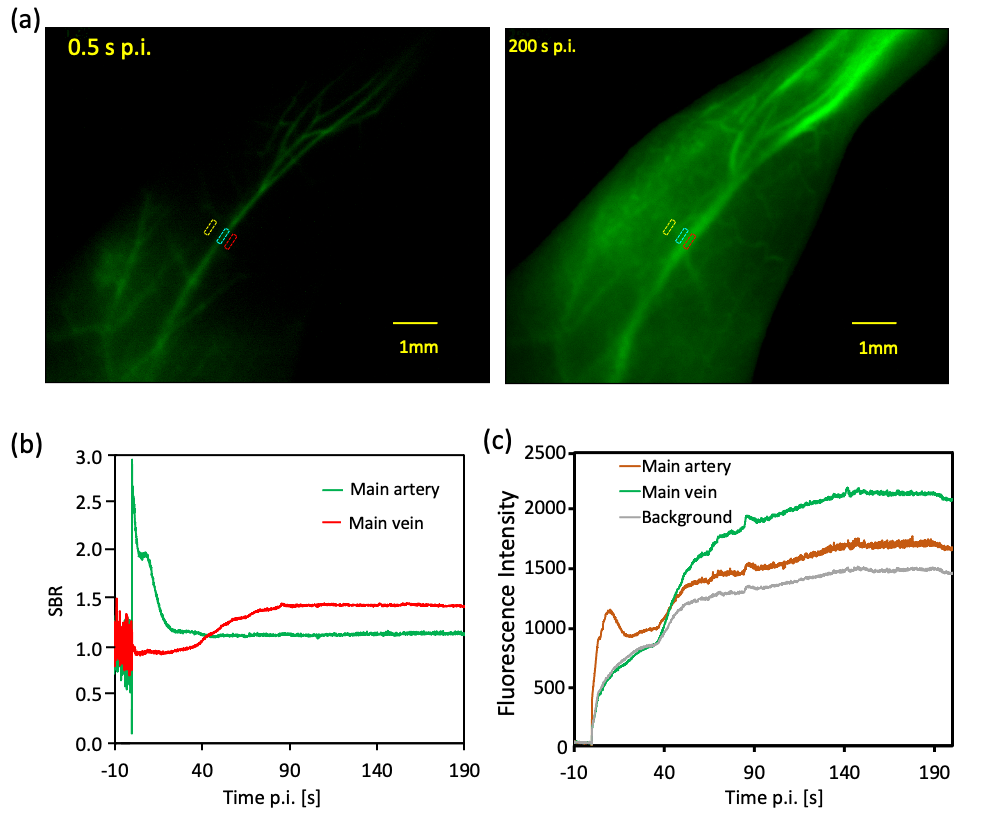

**Figure S10.** (a) The fluorescence imaging of hindlimb vessels (mouse lay on back) at 0.5 s and 200 s p.i. (post injection, the time zero is defined as the first observation of fluorescence in an artery) with a 1200 nm long pass filter, exposure time 50 ms, frame rate about 16 fps, excitation at 980 nm, 75 mW/cm^2^. (b) The signal to background ratio (SBR) as a function of time p.i., the region for artery signal indicated as cyanine rectangle, the region for vein signal indicated as red rectangle and the region for background signal indicated as yellow rectangle, as shown in (a). (c) The signal intensity as a function of time p.i., the region for artery signal indicated as cyanine rectangle, the region for vein signal indicated as red rectangle and the region for background signal indicated as yellow rectangle as shown in (a).

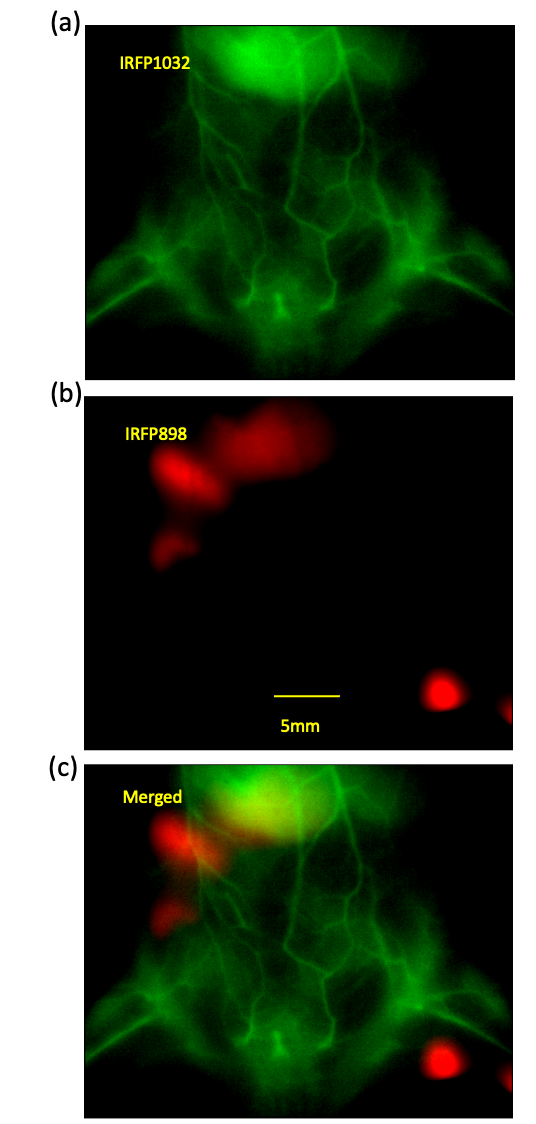

**Figure S11.** The dual color imaging of blood vessels and lymphatic networks by IRFP1032 and IRFP898, respectively. (a) The in vivo image of blood vessels (mouse lay on back) via intravenous injection of IRFP1032, excited at 980 nm, laser power 25 mW/cm^2^, 1200 nm long pass filter, exposure time 100 ms. (b) The in vivo image of lymphatic systems via intradermally injection of IRFP898 in the left hind footpads (2 hours post injection), excited at 825 nm, laser power 25 mW/cm^2^, 900 nm long pass filter and 1000 nm short pass filter, exposure time 100 ms, the liver signal could be observed clearly. (c) The merged image of (a) and (b).

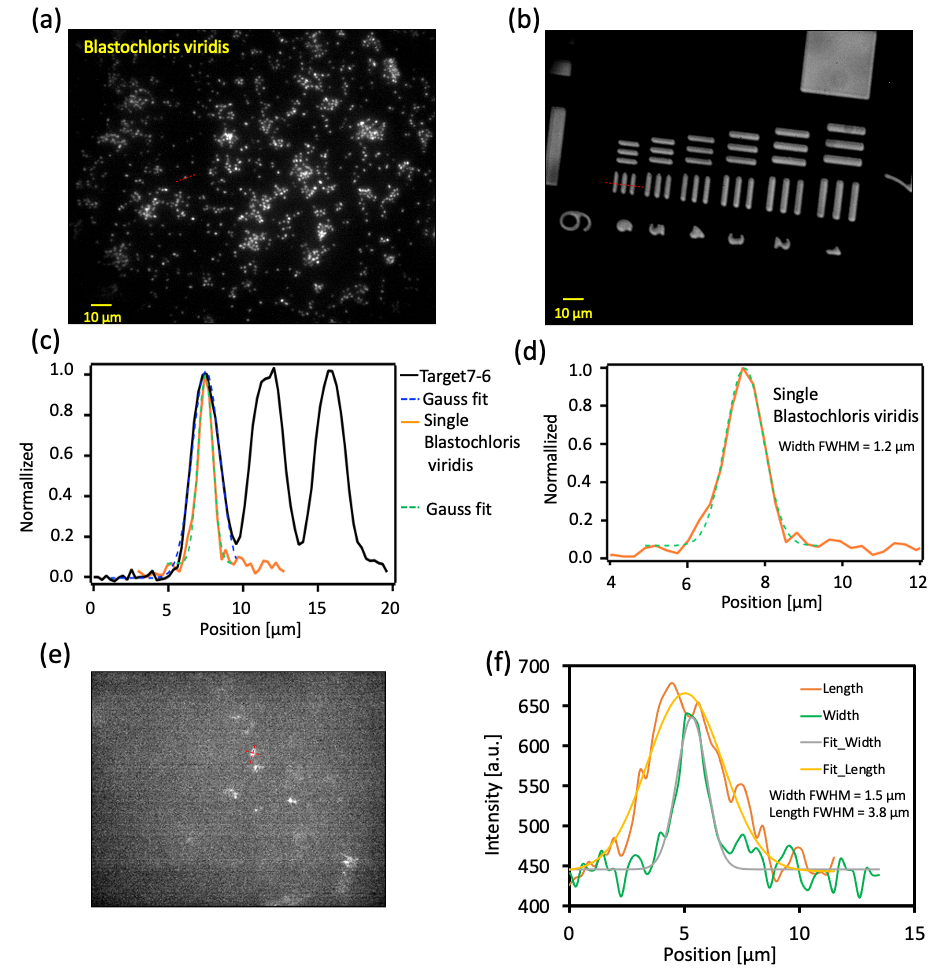

**Figure S12.** (a) The microscopy image of Blastochloris viridis on glass slide, objective 60x, excited at 980 nm, laser intensity 5 mW/cm^2^, 1100 nm long pass filter, exposure time 25 ms. (b) The microscopy image of 1951USAF resolution target, objective 60x, Tungsten Halogen lamp transmission image. (c) The intensity profile of the single Blastochloris viridis as indicated in (a) by red dash line, and the profile of the 6^th^ elements of the 7^th^ group of the 1951USAF resolution target, for which the line width is about 2.19 μm. (d) The gaussian fit of the single Blastochloris viridis with FWHM about 1.2 μm. (e) The bacteria clusters recorded in vivo, as shown in Figure 6c (0 ms). (f) The gaussian fit of the bacteria cluster of (e) indicated by a cross dotted red line, for length about 3.8 μm and the width about 1.5 μm.

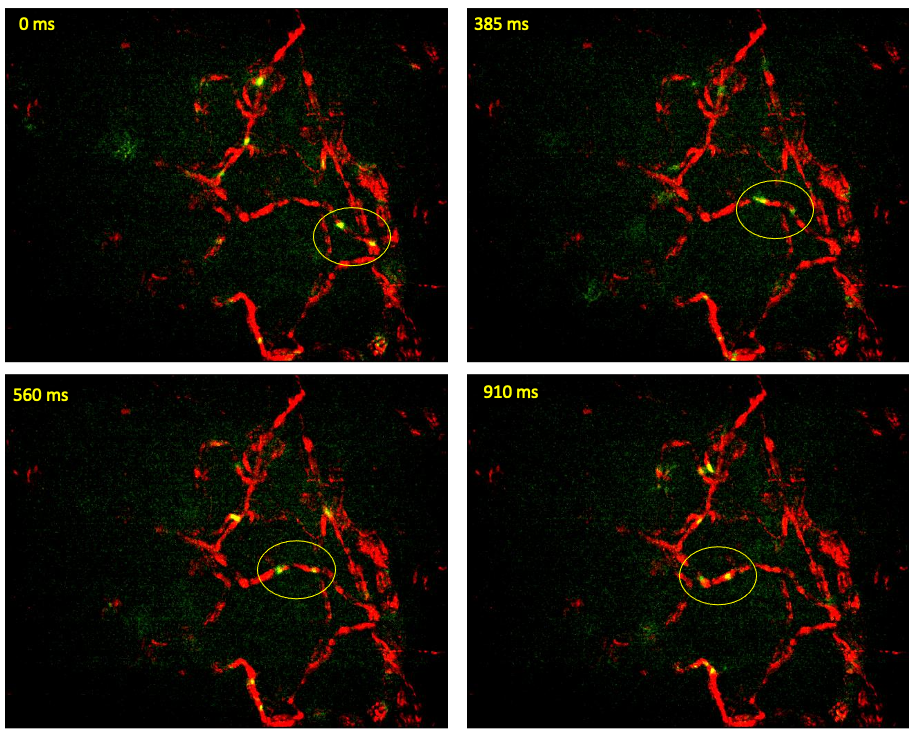

**Figure S13.** The in vivo tracking of a pair of bacteria clusters through the blood vessels of the mouse ear. We could find that the bacteria pair could be continuously recorded with similar distance in a relatively long period of time. There were no other bacteria that could be observed in this vessel during that time period. Therefore, the observations with frame rate about 30 fps is considered to be sufficient for tracking the bacteria in those capillaries, without frame missing, in the anaesthesia condition mouse.

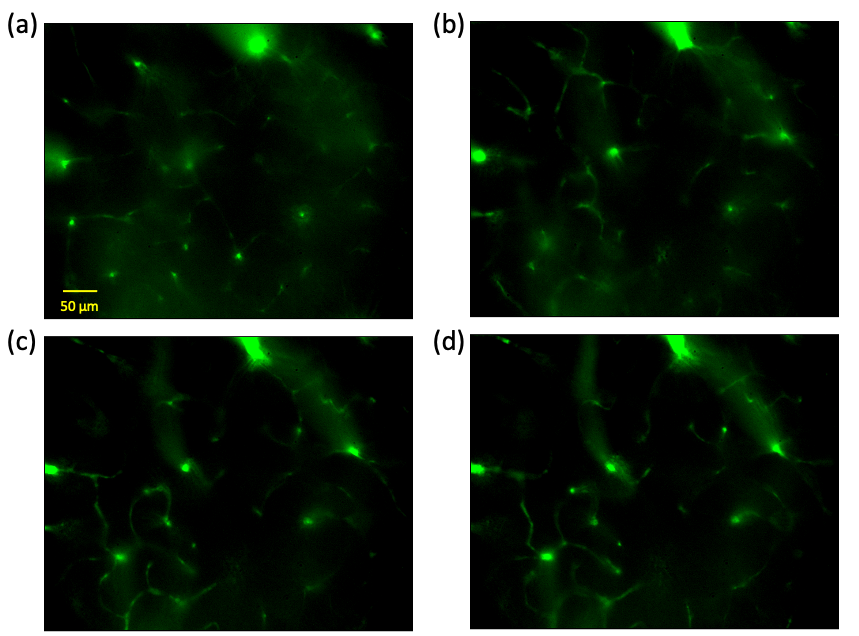

**Figure S14.** The in vivo microscopy image of brain blood vessels of mouse with cranial window via intravenous injection of IRFP1032, excited at 980 nm, laser power 25 mW/cm^2^, 1200 nm long pass filter, exposure time about 100 ms. (a), (b), (c), (d) were recorded at different focusing depths.

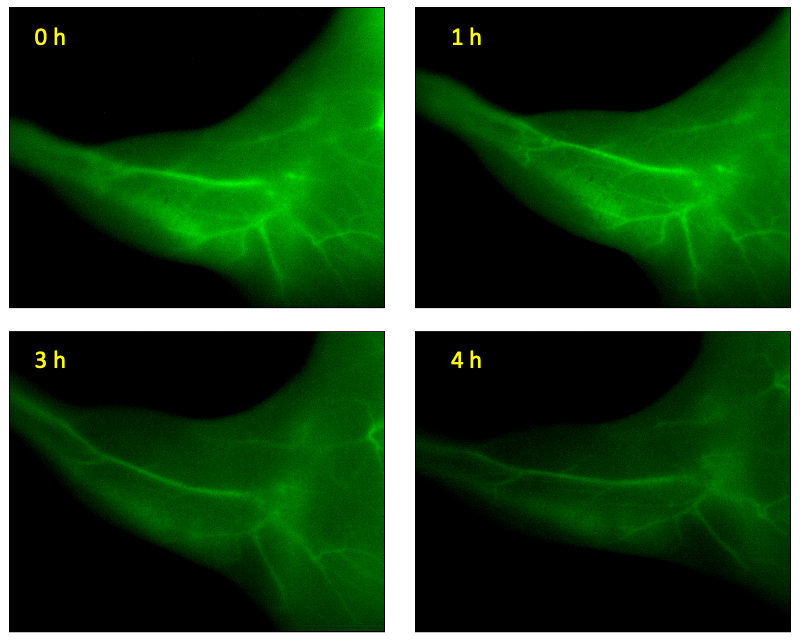

**Figure S15.** The fluorescence imaging of hindlimb vessels (mouse lay on back) at different time post injection, (0, 1, 3, 4 hours) with 1100 nm long pass filter, exposure time 100 ms, excitation at 980 nm, 25 mW/cm^2^. We could find that at 4 hours p.i. the blood vessels could be still visualized.

**Video 1.** Real-time in vivo blood flow dynamic imaging of hindlimb of a nude BALB/c mouse via intravenous injection of IRFP1032 with 1200 nm long pass filter, exposure time 50 ms, frame rate about 16 fps, excitation at 980 nm, 75 mW/cm^2^. The time zero is defined as the first sign of fluorescence in vessels.

**Video 2.** Real-time in vivo blood flow dynamic imaging of abdomen of a nude BALB/c mouse via intravenous injection of IRFP1032 with 1200 nm long pass filter, exposure time 100 ms, frame rate about 10 fps, excitation at 980 nm, 75 mW/cm^2^. The time zero is defined as the first sign of fluorescence in vessels.

**Video 3.** Real-time in vivo imaging of a nude BALB/c mouse via intravenous injection of IRFP1032 with 1200 nm long pass filter, exposure time 50 ms, frame rate about 16 fps, excitation at 980 nm, 75 mW/cm^2^ (green channel), and injection of IRFP898 in the hind footpads, excited at 825 nm, laser power 25 mW/cm^2^, 900 nm long pass filter and 1000 nm short pass filter, exposure time 50 ms, frame rate about 16 fps (red channel).

**Video 4.** Real-time in vivo imaging of single bacterial cell in the ear capillary of nude BALB/c mouse via intravenous injection of Blastochloris viridis with objective 60x, excited at 980 nm, laser intensity 75 mW/cm^2^, 1100 nm long pass filter, exposure time 25 ms, 30 fps.
